# Single-nucleus transcriptomic analysis reveals divergence of glial cells in peripheral somatosensory system between human and mouse

**DOI:** 10.1101/2022.02.15.480622

**Authors:** Donghang Zhang, Yiyong Wei, Jin Liu, Yaoxin Yang, Mengchan Ou, Yali Chen, Jiefei Shen, Tao Zhu, Cheng Zhou

**Author notes:** Address corresponding to: Dr. Cheng Zhou, Laboratory of Anesthesia and Critical Care Medicine, National-Local Joint Engineering Research Centre of Translational Medicine of Anesthesiology, West China Hospital of Sichuan University, Chengdu, 610041, China,; Dr. Jin Liu. Department of Anesthesiology, West China Hospital of Sichuan University, Chengdu, 610041, China,; Dr. Tao Zhu, Department of Anesthesiology, West China Hospital of Sichuan University, Chengdu, 610041, China,.

## Abstract

Glial cells play a crucial role in regulating physiological and pathological functions, such as sensation, infections, acute injuries, and chronic neurodegenerative disorders. Glial cells include astrocytes, microglia, and oligodendrocytes in the central nervous system (CNS) and satellite glial cells (SGCs) in the peripheral nervous system (PNS). Despite the understanding of glial subtypes and functional heterogeneity in animal models achieved by single-cell or single-nucleus RNA sequencing, no research has investigated the transcriptomic profiles of glial cells in the human PNS and spinal cord. Here, we used high-throughput single-nucleus RNA sequencing to map the cellular and molecular heterogeneity of SGCs in the human dorsal root ganglion (DRG) and astrocytes, microglia, and oligodendrocytes in the human spinal cord. To explore the conservation and divergence across species, we compared these human findings with those from mice. Additionally, the expression profiles of risk genes of common DRG and spinal cord diseases in glial cells were compared between humans and mice. As a result, little SGCs heterogeneity was found in both human and mouse DRG. In the human spinal cord, astrocytes, microglia, and oligodendrocytes were respectively divided into six distinct transcriptomic subclusters. In the mouse spinal cord, astrocytes, microglia, and oligodendrocytes were divided into six, five, and six distinct transcriptomic subclusters, respectively. The comparative results revealed substantial heterogeneity in all glial cells between humans and mice. Notably, we also identified transcriptomic heterogeneity in several classical genes and risk genes for neurological disorders across humans and mice. Together, the present data comprehensively profiled glial cell heterogeneity and provides a powerful resource for investigating the cellular basis of glial-related physiological and pathological conditions in peripheral somatosensory system.

## Introduction

In addition to neurons, both the peripheral (PNS) and central nervous system (CNS) include glial cells. The major types of glial cells in the CNS include astrocytes, microglia, and oligodendrocytes [1, 2], whereas in the PNS, satellite glial cells (SGCs) are the major glial cells, which are the counterpart of astrocytes [3, 4]. Astrocytes are the most abundant glial cell in the mammalian CNS and play a pivotal role from structural support of the CNS to regulation of neuroinflammation associated with many neurological and psychiatric diseases [5]. Microglia are well known as the primary resident immune cells of the CNS, providing the initial host defense and orchestrating immune responses [6]. Oligodendrocytes are responsible for axon myelination and enabling fast saltatory impulse propagation within the CNS, and associated with many neurodegenerative diseases [7, 8].

Despite the clear evidence that glial cells of peripheral somatosensory system, including dorsal root ganglion (DRG) and spinal cord, play a pivotal role in sensory perception, processing as well as motor behavior, their cellular and functional heterogeneity remain unclear. Recently, emerging evidence has revealed cellular and molecular heterogeneity in specific glial cells of the CNS using single-cell (scRNA-seq) or single-nucleus RNA sequencing (snRNA-seq) [3, 9–12]. Using scRNA-seq, Hasel et al. identified 10 astrocyte subclusters in the mouse brain and characterized the heterogeneous responses of different astrocyte subsets to inflammation [12]. van Weperen et al. performed scRNA-seq on mouse stellate ganglia and divided SGCs into six distinct transcriptomic subtypes [3]. Zheng et al. performed scRNA-seq to compare microglial subtypes in the cortex and spinal cord identifying three subclusters in the cortex and two in the spinal cord and showing distinct subpopulations between spinal and cortical microglia [13]. Using a combination of scRNA-seq and multicolor flow cytometry, Sousa et al. identified distinct microglial activation profiles under inflammation, which are substantially different from neurodegenerative disease-related profiles [14]. Using scRNA-seq, mature oligodendrocytes could be classified into six distinct subpopulations, which show different responses to spinal cord injury, in the mouse CNS [10, 15]. Although the cellular and functional heterogeneity of glial cells in the mouse nervous system is relatively well known, limited information is available for humans, particularly about those in the peripheral DRG and spinal cord.

In this study, we used 10× Genomics snRNA-seq to map the heterogeneity of glial cells, including SGCs, astrocytes, microglia, and oligodendrocytes, in the human DRG and spinal cord. Additionally, we addressed interspecies heterogeneity by comparing human and mouse transcriptomic profiles. This study aimed to serve as an important resource for future research on the molecular basis of human physiological and pathological conditions related to glial cells.

## Results

### Identification of glial cell types in human and mouse DRG

snRNA-seq was performed on 26,279 nuclei from the L3-L5 DRG of three adult donors (Figure 1A). DRG cells were initially classified into 11 clusters based on their distinct transcriptional characteristics (Figure 1B). SGCs (10.7% of total nuclei) were labeled with the representative markers *FABP7* and *APOE* and formed Cluster 2 (Figure 1C-D, Supplementary Figure 1A). We also reanalyzed available snRNA-seq transcriptional data from the mouse DRG from a previous study [16]. DRG cells from seven C57 mice were classified into 22 cell types (Figure 1E), of which Cluster 0 was identified as SGCs (9.8% of total nuclei) through the markers *Fabp7* and *Apoe* (Figure 1F and G, Supplementary Figure 1B).

**Figure 1.**
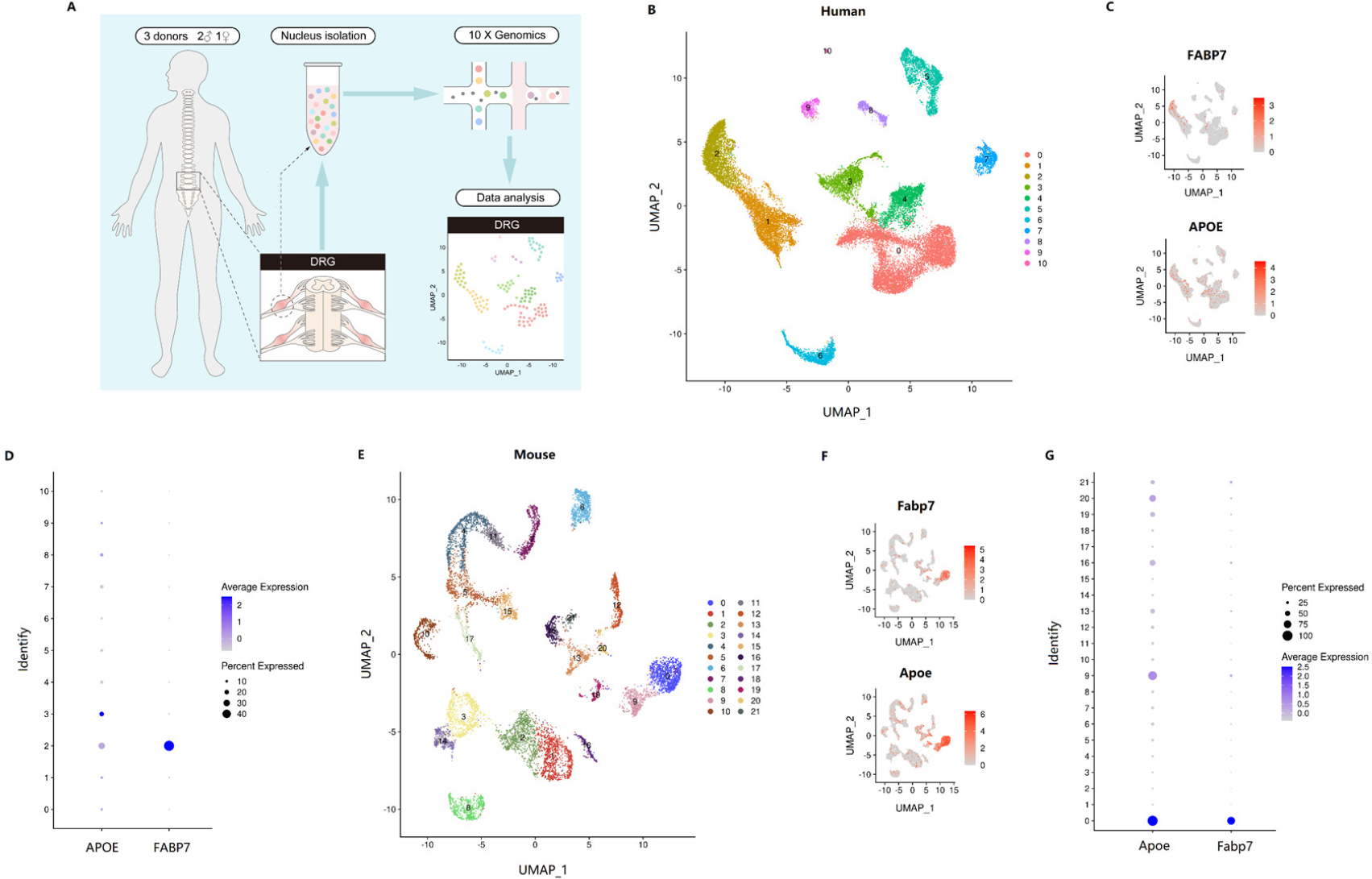
Identification of glial cell-types in DRG. (A) Overview of the experimental workflow for snRNA-seq in DRG. (B) UMAP plot of human DRG cells showing 11 major cell types. Dots, individual cells; colors, cell types. (C) UMAP plot showing the expression of representative well-known marker genes of human SGCs. Numbers reflect the number of unique molecular identifiers (UMIs) detected for the specified gene in each cell. (D) Dot plot showing the distribution of expression levels of human SGCs marker genes across all cell types. (E) UMAP plot of mouse DRG cells showing 22 major cell types. (F) UMAP plot showing the expression of representative well-known marker genes of mouse SGCs. (G) Dot plot showing the distribution of expression levels of mouse SGCs marker genes across all cell types. UMAP, Uniform Manifold Approximation and Projection; SGCs, satellite glial cells.

### Identification of cell types in human and mouse spinal cord

The lumbar enlargements of spinal cords of the same three adult donors were also extracted for snRNA-seq (Figure 2A). As a result, 28,677 nuclei were isolated and initially classified into eight clusters (Figure 2B). Astrocytes (9.6% of total nuclei) were identified with the representative markers *ATP1A2*, *AQP4*, *GJA1*, and *SLC1A2* as Cluster 2 (Figure 2C and F, Supplementary Figure 1C). Microglia (17.6% of total nuclei) were identified with *PTPRC*, *CTSS*, and *ITGAM* (Figure 2D and F, Supplementary Figure 1C) as Cluster 1, and oligodendrocytes (52.4% of total nuclei) were identified as Cluster 0 with *MBP*, *MOBP*, *MOG*, and *PLP1* (Figure 2E and F, Supplementary Figure 1C). For the snRNA-seq analysis of the mouse spinal cord, we extracted transcriptional data of astrocytes (8.6%), microglia (1.2%), and oligodendrocytes (15.4%) from a previous study [17].

**Figure 2.**
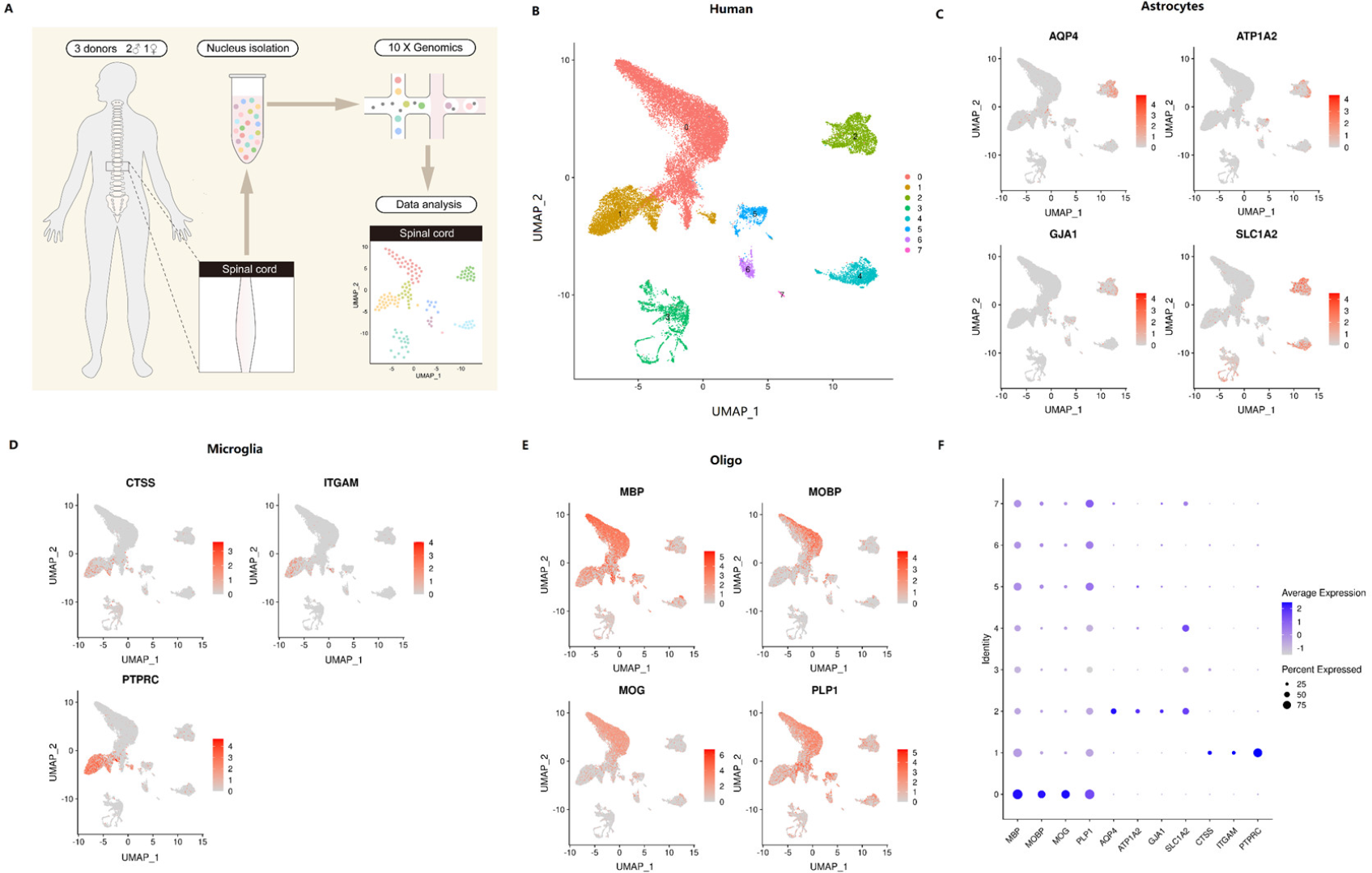
Identification of glial cell-types in spinal cord. (A) Overview of the experimental workflow for snRNA-seq in human spinal cord. (B) UMAP plot of human spinal cells showing 8 major cell types. Dots, individual cells; colors, cell types. (C-E) UMAP plot showing the expression of representative well-known marker genes of human spinal astrocytes (C), microglia (D), and oligodendrocytes (E). Numbers reflect the number of unique molecular identifiers (UMIs) detected for the specified gene in each cell. (F) Dot plot showing the distribution of expression levels of marker genes of spinal astrocytes, microglia, and oligodendrocytes across all cell types. UMAP, Uniform Manifold Approximation and Projection.

### Human-mouse divergence

To address the similarities and differences in glial cells between humans and mice, we performed co-clustering methods [18] to align human transcriptomic data with those from mice (Figure 3). Alignment results revealed significant heterogeneity in all types of glial cells between mice and humans, including SGCs (Figure 3A), astrocytes (Figure 3B), microglia (Figure 3C), and oligodendrocytes (Figure 3D), as indicated by the slight overlap across species.

**Figure 3.**
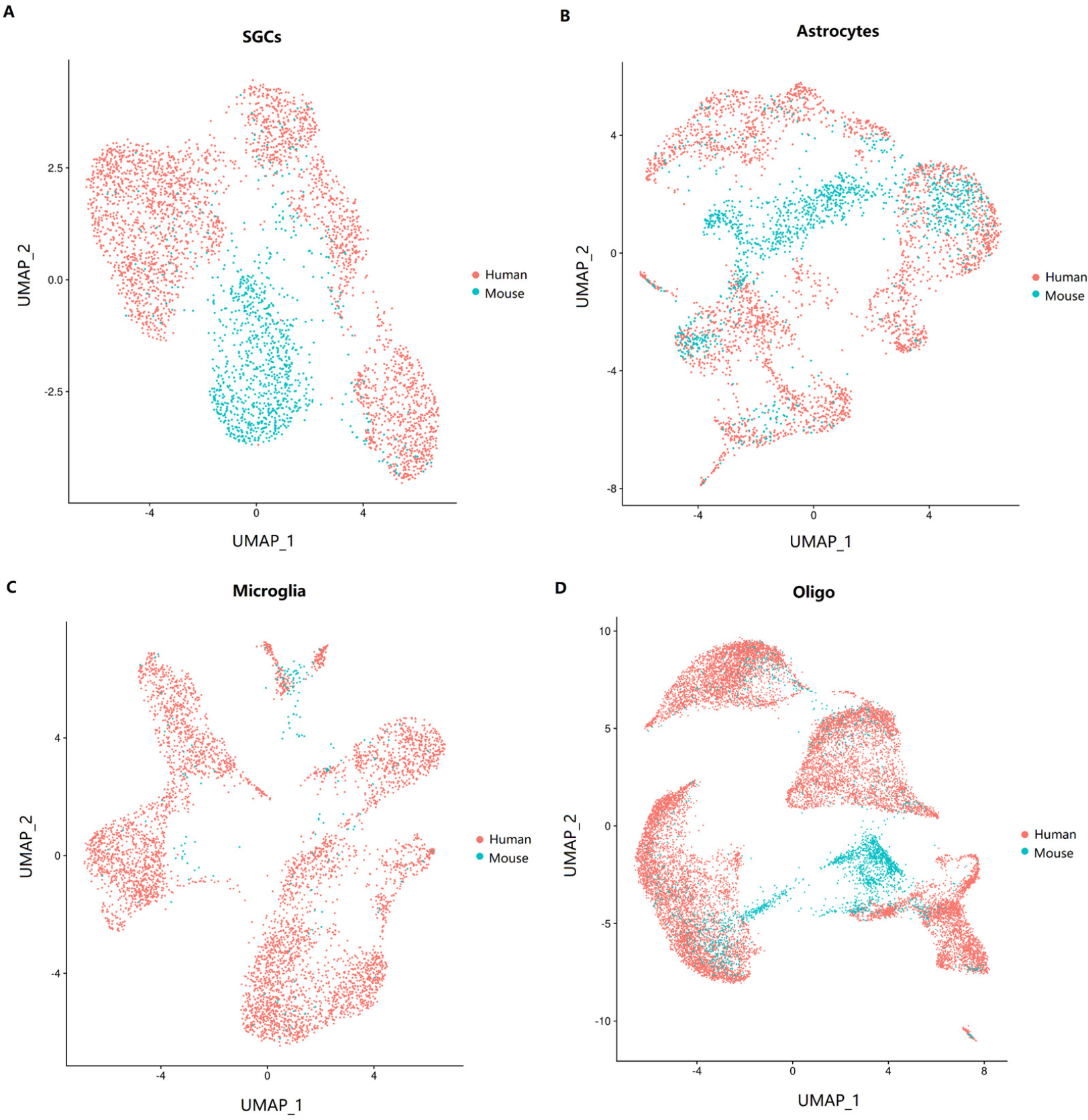
Divergence of glial cells across human-mouse. (A-D) UMAP plot showing the co-clustering of SGCs (A), astrocytes (B), microglia (C), and oligodendrocytes (D) between human and mouse. UMAP, Uniform Manifold Approximation and Projection; SGCs, satellite glial cells; oligo, oligodendrocytes.

### Identification of glial cell subtypes

To determine the heterogeneity within each glial cell type, SGCs were further classified into six subclusters in humans (Figure 4A and B, Supplementary Figure 2A) and three subclusters in mice (Figure 4C and D, Supplementary Figure 2B). Regarding intraspecies comparison, DRG SGCs showed substantial similarity across subtypes, as they could not be classified completely. In human spinal cord, we classified astrocytes (Figure 5A and B, Supplementary Figure 3A), microglia (Figure 5C and D, Supplementary Figure 3B), and oligodendrocytes (Figure 5E and F, Supplementary Figure 3C) into six subclusters, respectively, based on their transcriptional characteristics. And in mouse spinal cord, we classified astrocytes (Figure 6A and B, Supplementary Figure 3D), microglia (Figure 6C and D, Supplementary Figure 3E), and oligodendrocytes (Figure 6E and F, Supplementary Figure 3F) into six, five, and six subclusters, respectively. We further analyzed the representative marker genes which exhibited uniquely high expression levels in each cluster, and finally identified several marker genes that associated with somatosensory system disorders (Figure 5B, D, F and Figure 6B, D, and F), such as *LGR6* (leucine rich repeat containing G protein-coupled receptor 6) in Cluster 2 of human astrocytes, *Gfap* (glial fibrillary acidic protein) and *Aqp4* (aquaporin 4) in Cluster 2 of mouse astrocytes, *EZH2* (enhancer of zeste 2 polycomb repressive complex 2 subunit) in Cluster 4 of human microglia, *Ctss* (cathepsin S), *Apoe* (apolipoprotein E), and *Hpgds* (hematopoietic prostaglandin D synthase) in Cluster 1 of mouse microglia, *S100a9* (S100 calcium binding protein A9) and *Anxa1* (annexin A1) in Cluster 4 of mouse microglia, and *CACNA1B* (calcium voltage-gated channel subunit alpha1 B) in Cluster 3 of human oligodendrocytes.

**Figure 4.**
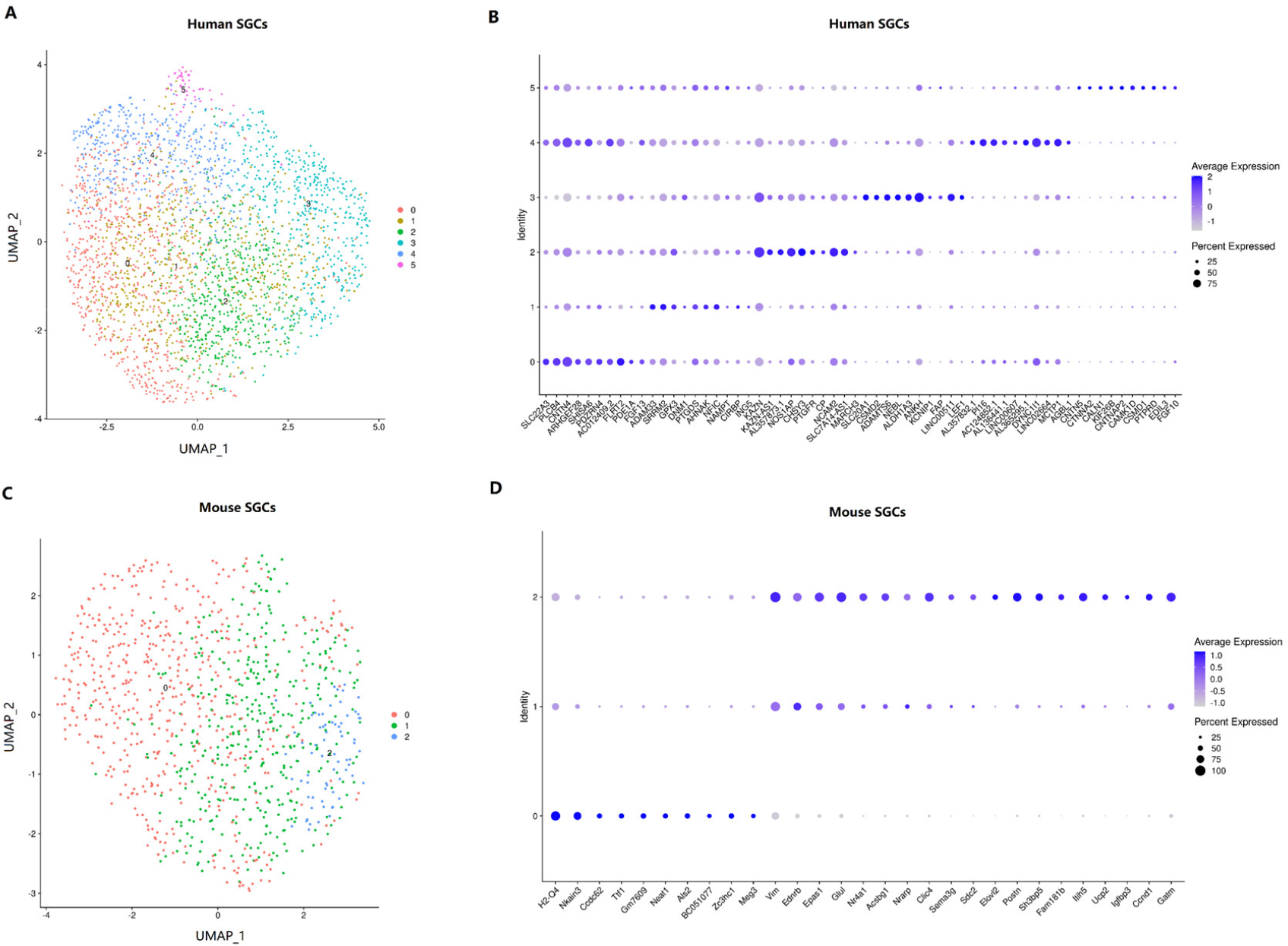
Identification of subtypes of DRG SGCs. (A) UMAP plot showing 6 clusters in human SGCs. Dots, individual cells; colors, SGCs clusters. (B) Dot plot showing the expression of the top ten most differentially expressed genes across all the SGCs clusters in humans. (C) UMAP plot showing 3 clusters in mouse SGCs. (D) Dot plot showing the expression of the top ten most differentially expressed genes across all the SGCs clusters in mice. UMAP, Uniform Manifold Approximation and Projection; SGCs, satellite glial cells.

**Figure 5.**
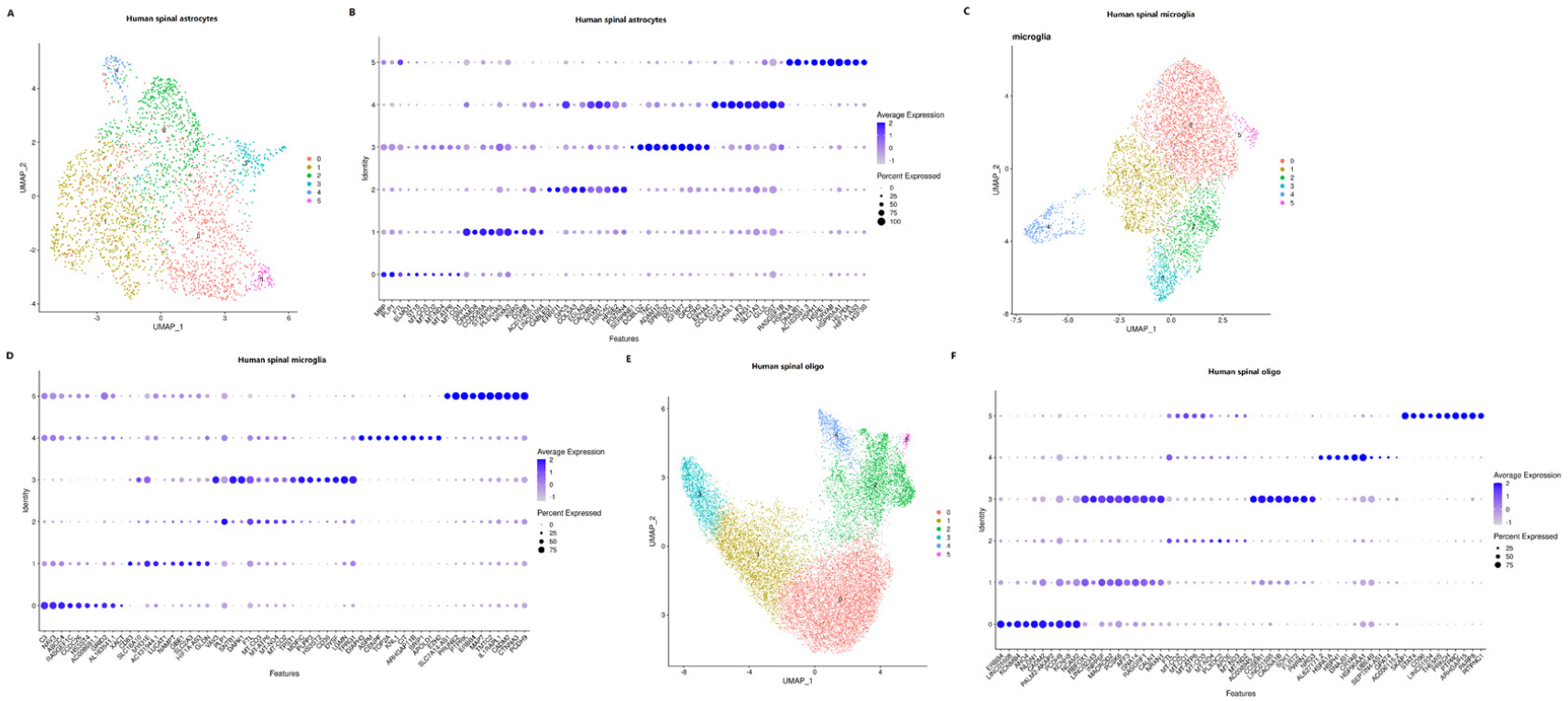
Identification of subtypes of astrocytes, microglia, and oligodendrocytes in humans. (A, C, E) UMAP plot showing subclusters of astrocytes (A), microglia (C), and oligodendrocytes (E) in human spinal cord. Dots, individual cells. (B, D, F) Dot plot showing the expression of the top ten most differentially expressed genes across all the astrocytes (B), microglia (D), and oligodendrocytes (F) clusters in humans. UMAP, Uniform Manifold Approximation and Projection; oligo, oligodendrocytes.

**Figure 6.**
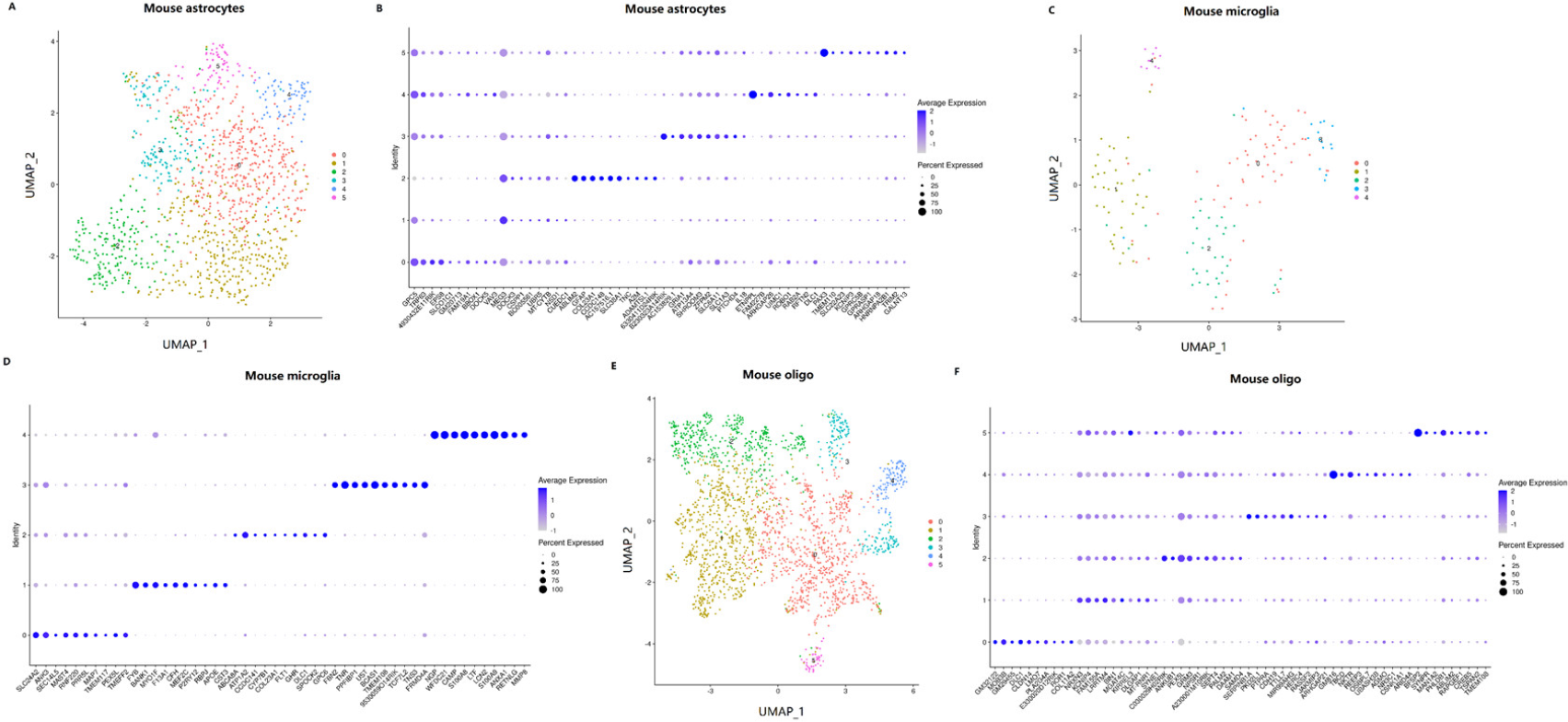
Identification of subtypes of astrocytes, microglia, and oligodendrocytes in mice. (A, C, E) UMAP plot showing subclusters of astrocytes (A), microglia (C), and oligodendrocytes (E) in mouse spinal cord. Dots, individual cells. (B, D, F) Dot plot showing the expression of the top ten most differentially expressed genes across all the astrocytes (B), microglia (D), and oligodendrocytes (F) clusters in mouse spinal cord. UMAP, Uniform Manifold Approximation and Projection; oligo, oligodendrocytes.

### Transcriptional profiles of classical markers

Transcriptional profiles of classical markers for somatosensory system in glial cells between humans and mice were compared, including ion channels, neurotransmitter receptors, neuropeptides, and transcription factors (Figures 7–10).

**Figure 7.**
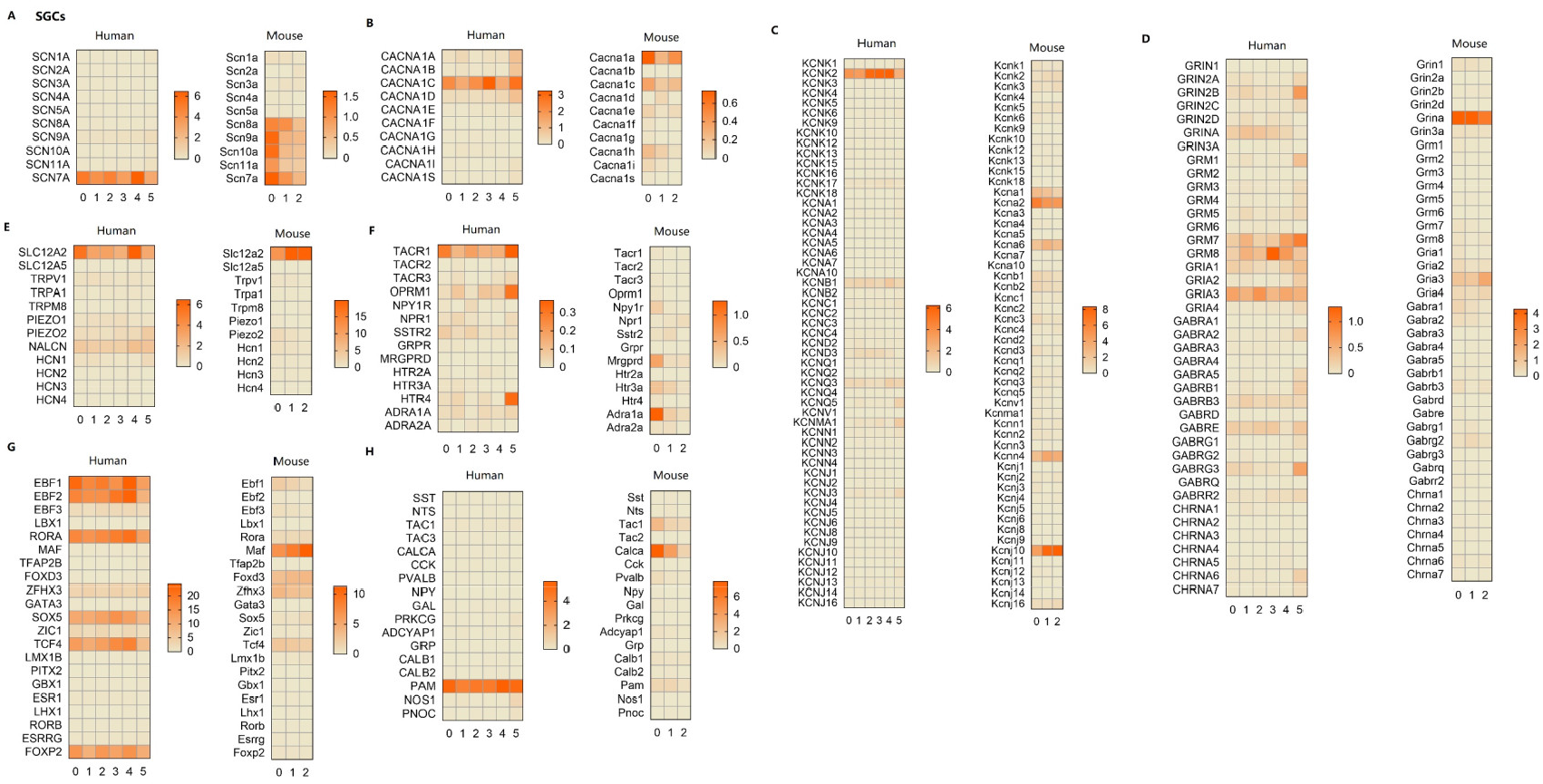
Expressional profiles of classical marker genes in DRG SGCs. Normalized mean gene expression of classic marker genes of sodium channels (A), calcium channels (B), potassium channels (C), glutamatergic-, GABAergic-, and cholinergic receptor (D), other ion channels (E), other receptors (F), transcription factors (G), and neuropeptides (H) in human and mouse SGCs. SGCs, satellite glial cells.

**Figure 8.**
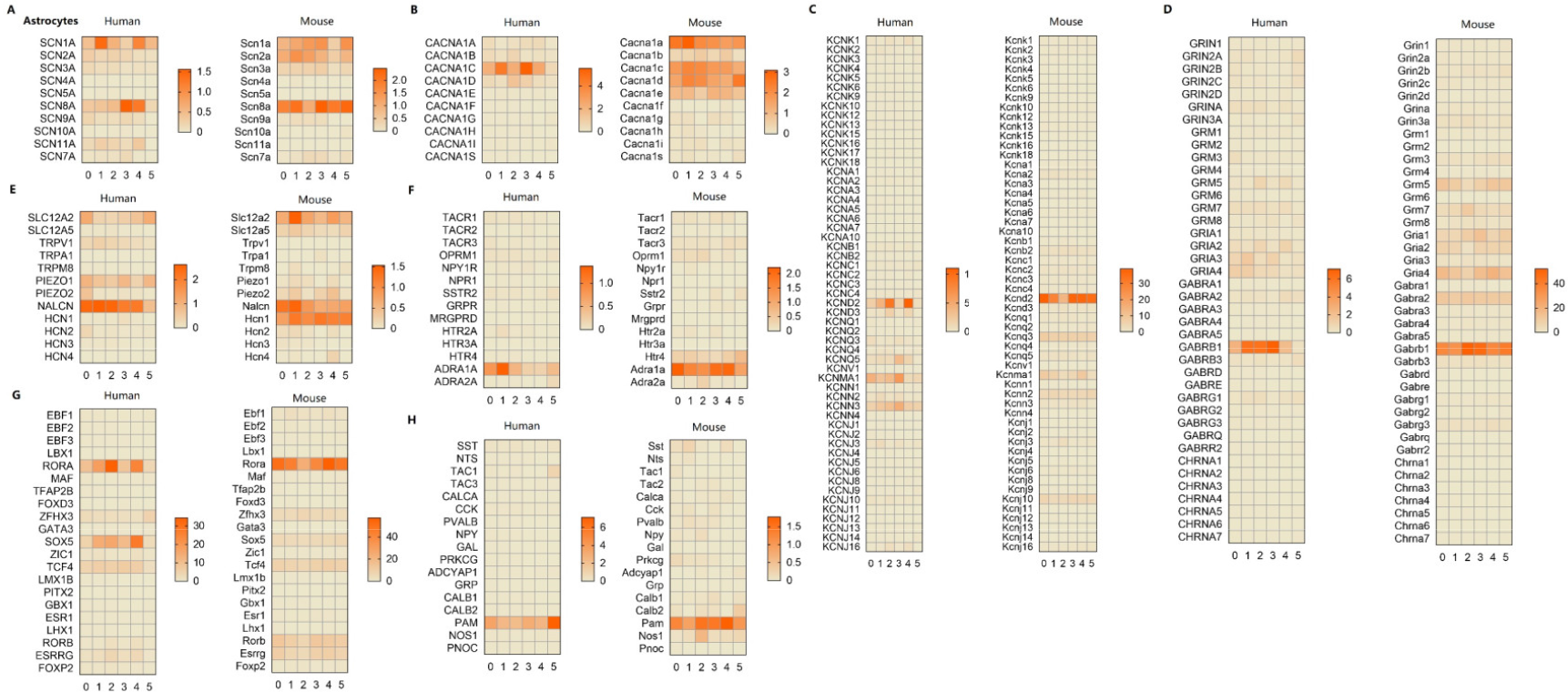
Expressional profiles of classical marker genes in spinal astrocytes. Normalized mean gene expression of classic marker genes of sodium channels (A), calcium channels (B), potassium channels (C), glutamatergic-, GABAergic-, and cholinergic receptor (D), other ion channels (E), other receptors (F), transcription factors (G), and neuropeptides (H) in human and mouse astrocytes.

**Figure 9.**
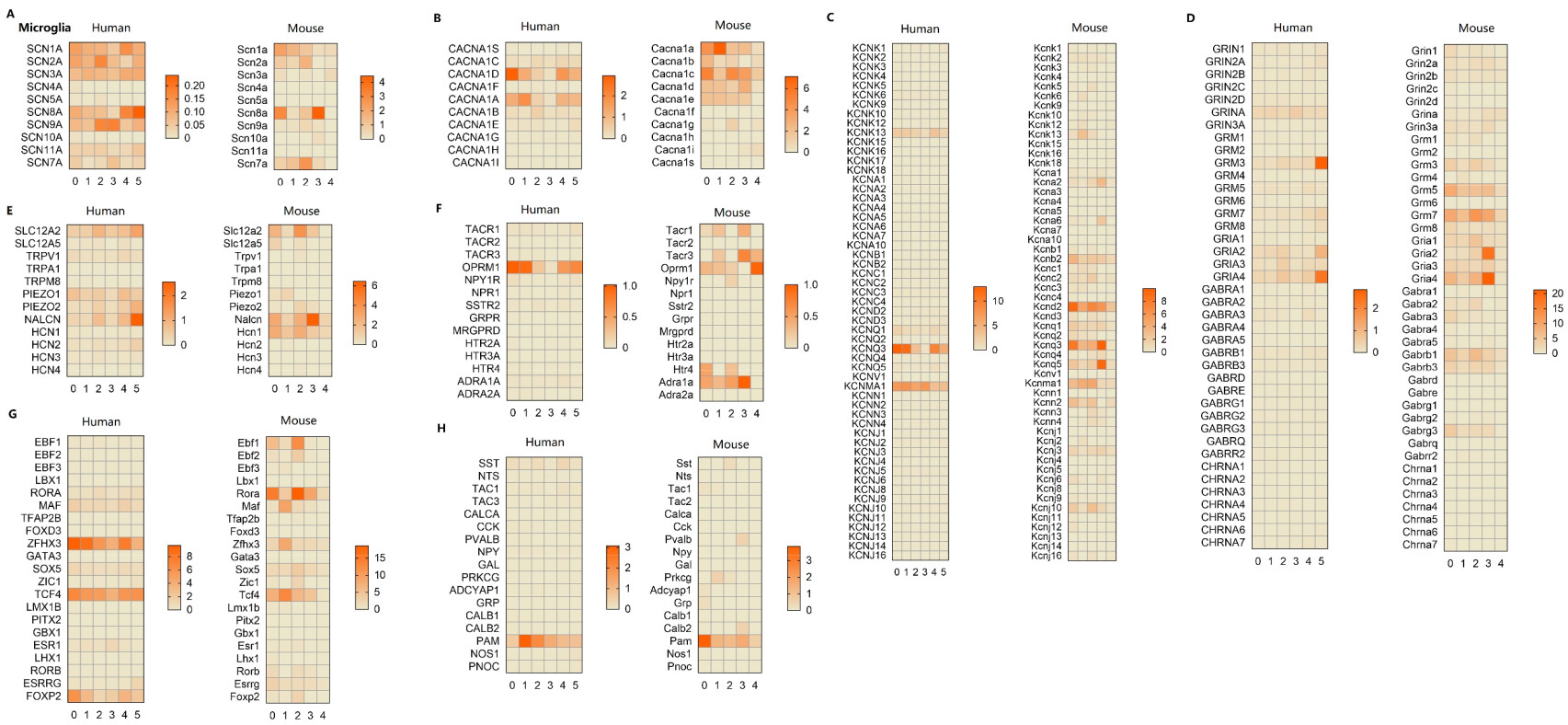
Expressional profiles of classical marker genes in microglia. Normalized mean gene expression of classic marker genes of sodium channels (A), calcium channels (B), potassium channels (C), glutamatergic-, GABAergic-, and cholinergic receptor (D), other ion channels (E), other receptors (F), transcription factors (G), and neuropeptides (H) in human and mouse microglia.

**Figure 10.**
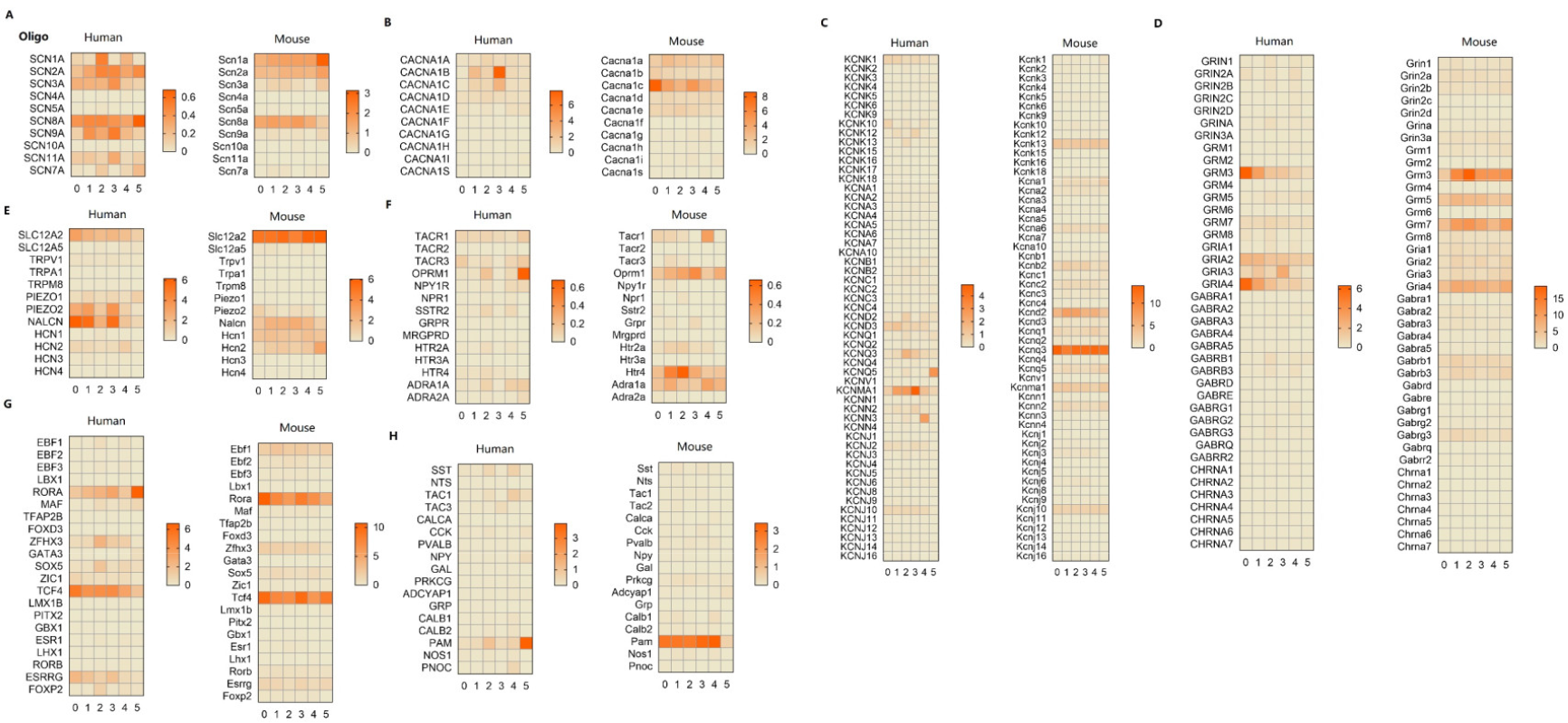
Expressional profiles of classical marker genes in oligodendrocytes. Normalized mean gene expression of classic marker genes of sodium channels (A), calcium channels (B), potassium channels (C), glutamatergic-, GABAergic-, and cholinergic receptor (D), other ion channels (E), other receptors (F), transcription factors (G), and neuropeptides (H) in human and mouse oligodendrocytes.

First, substantial similarity in marker expression was found across SGCs intra-clusters (Figure 7). For example, the expression levels of most ion channels, including voltage-gated sodium channels, calcium channels, and potassium channels, were comparable across subclusters in human and mouse DRG SGCs (Figure 7A-C). Nevertheless, several genes were differentially expressed across clusters. For example, *HTR4* (5-hydroxytryptamine receptor 4, which modulates the release of various neurotransmitters in both PNS and CNS) showed significantly higher expression levels in Cluster 5 than in other clusters in human DRG SGCs (Figure 7F, left panel). *Adra1a* (Adrenoceptor alpha 1A, which regulates cell growth and proliferation) showed significantly higher expression levels in Cluster 0 than in other clusters in mouse DRG SGCs (Figure 7F, right panel).

In astrocytes, microglia, and oligodendrocytes, the expression levels of multiple genes showed significant heterogeneity across subclusters, such as *SCN8A* (Na_v_1.6, essential for rapid membrane depolarization during action potential formation in excitable cells) (Figure 8A), *KCND2* (K_v_4.2, which mediates a rapidly inactivating response in neurons) (Figure 8C), *scn8a* (Figure 9A), *CACNA1D* (Ca_v_1.3, which mediates Ca^2+^ entry into excitable cells) (Figure 9B), *GRM3* (glutamate metabotropic receptor 3, which is associated with glutamatergic neurotransmission) (Figure 9D), *NALCN* (sodium leak channel, which conducts a persistent sodium leak current contributing to tonic neuronal excitability) (Figure 9E), *Tacr3* (tachykinin receptor 3) (Figure 9F), *Maf* (MAF bZIP transcription factor, which regulates various biological processes, such as apoptosis) (Figure 9G), *CACNA1B* (Ca_v_2.2, which controls neuron neurotransmitter release) (Figure 10B), *Cacna1c* (Ca_v_1.2, which mediates inward Ca^2+^ influx upon membrane polarization) (Figure 10B), *KCNQ5* (K_v_7.5, which yields currents that activate slowly with depolarization) (Figure 10C), *KCNN3* (KCa2.3, which regulates neuronal excitability by contributing to the slow component of synaptic afterhyperpolarization) (Figure 10C), *OPRM1* (opioid receptor mu 1, which modulates analgesia and drug dependence) (Figure 10F), *Tacr1* (tachykinin receptor 1, which encodes the receptor for tachykinin substance P) (Figure 10F), and *PAM* (peptidylglycine alpha-amidating monooxygenase, which catalyzes the conversion of neuroendocrine peptides to active alpha-amidated products) (Figure 10H).

Next, we compared the transcriptomic data of these marker genes in glial cells between humans and mice. The overall transcriptional patterns showed a significant divergence between humans and mice. For example, *Scn9a* (Na_v_1.7, which mediates nociception signaling) (Figure 7A), *Scn10a* (Na_v_1.8, involved in the onset of pain associated with peripheral neuropathy) (Figure 7A), *Calca* (calcitonin gene-related peptide CGRP, which functions as a vasodilator) (Figure 7B), *Kcnj10* (KIR4.1, responsible for the potassium buffering action of glial cells) (Figure 7C), *Kcna2* (K_v_1.2, which allows nerve cells to efficiently repolarize following an action potential) (Figure 7C), and *Maf* (Figure 7G) showed high expression in mouse DRG SGCs, but slightly in humans. *Cacna1a* (Figure 8B) and *Hcn1* (hyperpolarization activated cyclic nucleotide gated potassium channel 1, which contributes to the native pacemaker currents in neurons) (Figure 8E) was highly expressed in mouse astrocytes, but slightly in humans. *Cacna1a* (Figure 9B), *Kcnd2* (Figure 9C), *Hcn1* (Figure 9E), *Tacr1* (Figure 9F), *Htr4* (Figure 9F), *Adra1a* (Figure 9F), and *Rora* (RAR-related orphan receptor A, which functions as a nuclear hormone receptor) (Figure 9G) were highly expressed in mouse microglia, but slightly in humans. *Hcn1* (Figure 10E), *Kcnq3* (Figure 10C), and *Grm5* (glutamate metabotropic receptor 5, involved in the regulation of neural network activity and synaptic plasticity) (Figure 10D) were highly expressed in mouse oligodendrocytes but slightly in humans.

### Disease-associated genes in glial cells

To better understand the conservation and divergence of genes associated with DRG and spinal cord diseases between humans and mice, we compared the transcriptional profiles of these risk genes. First, we examined gene expression levels in two types of chronic pain, namely, neuropathic and inflammatory pain (Figures 11-12). Several neuropathic pain-related genes, including *MAPK1/Mapk1* (mitogen-activated protein kinase 1, which acts as an integration point for multiple biochemical signals), *HMGB1/Hmgb1* (high mobility group box 1, which plays a role in inflammation process), and *STAT3/stat3* (signal transducer and activator of transcription 3, which mediates the expression of various genes in response to cell stimuli) showed high expression levels in both humans (Figure 11A, left panel) and mice (Figure 11A, right panel). Notably, *NGF/Ngf* (nerve growth factor, involved in the regulation of growth and differentiation of sympathetic and certain sensory neurons) was highly expressed in human SGCs but slightly in mouse SGCs (Figure 11A). *CALCA/Calca*, *GJA1/Gja1* (gap junction protein alpha 1, associated with cell– cell gap junctions), and *CXCR4/Cxcr4* (C-X-C motif chemokine receptor 4, associated with pain signaling) were highly expressed in mouse SGCs but slightly in humans (Figure 11A). The inflammatory pain-related gene *PRKCA* (protein kinase C alpha, which modulates different cellular processes, such as cell adhesion and cell transformation) showed extremely high expression levels in human SGCs (Figure 11B) but slightly in mice, whereas *MAPK3/Mapk3* (mitogen-activated protein kinase 3) was highly expressed in mouse SGCs but slightly in humans (Figure 11B).

**Figure 11.**
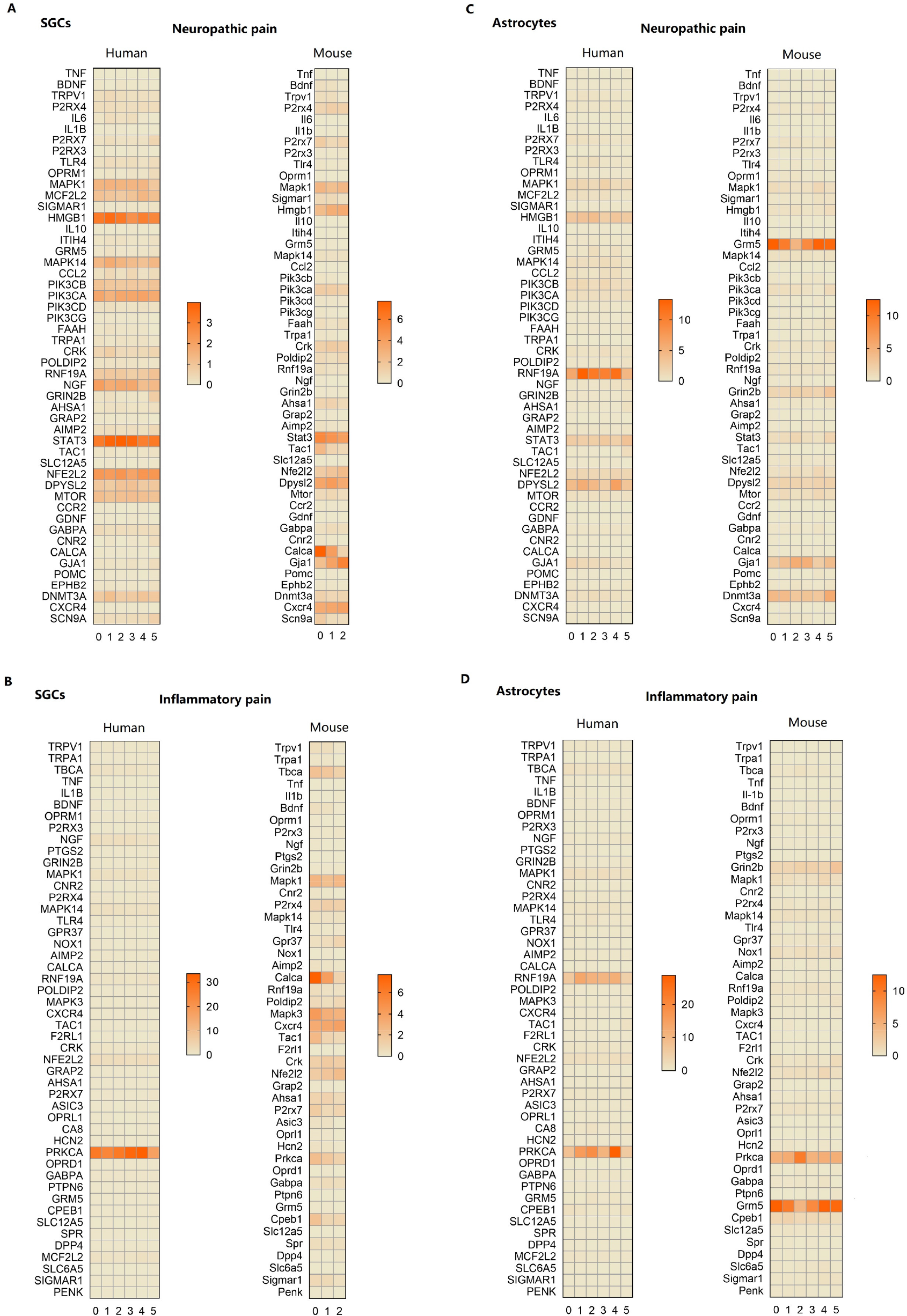
Expressional profiles of disease-risk genes in SGCs and astrocytes. (A-B) The transcriptional profiles of risk genes for neuropathic pain (A) and inflammatory pain (B) in human and mouse SGCs. (C-D) The transcriptional profiles of risk genes for neuropathic pain (C) and inflammatory pain (D) in human and mouse astrocytes.

**Figure 12.**
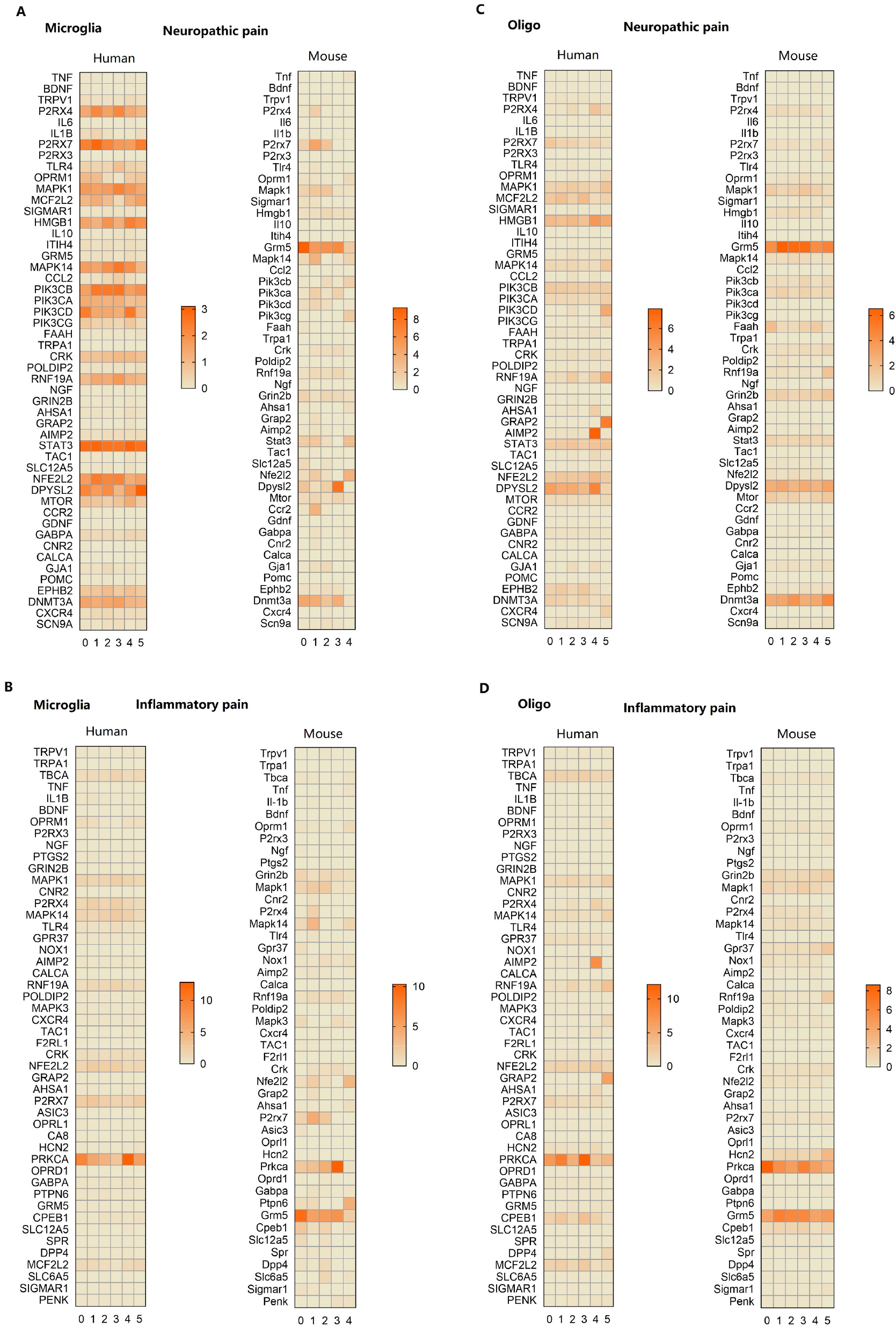
Expressional profiles of disease-risk genes in microglia and oligodendrocytes. (A-B) The transcriptional profiles of risk genes for neuropathic pain (A) and inflammatory pain (B) in human and mouse microglia. (C-D) The transcriptional profiles of risk genes for neuropathic pain (C) and inflammatory pain (D) in human and mouse oligodendrocytes.

In astrocytes, the common neuropathic/inflammatory pain genes *RNF19A/Rnf19a* (ring finger protein 19A, RBR E3 ubiquitin protein ligase, involved in amyotrophic lateral sclerosis [ALS] and Parkinson’s disease) were highly expressed in humans but not in mice, whereas *GRM5/Grm5* was highly expressed in mice, but slightly in humans (Figure 11C and D). Neuropathic pain-related genes *HMGB1/Hmgb1* and *DPYSL2/Dpysl2* (dihydropyrimidinase-like 2, which regulates synaptic signaling through interactions with calcium channels) were highly expressed in human astrocytes, but slightly in mice (Figure 11C). *DNMT3A/Dnmt3a* (DNA methyltransferase 3 alpha, which functions in de novo methylation) was highly expressed in mouse astrocytes but slightly in humans (Figure 11C). On the other hand, the inflammatory pain-related genes *PRKCA/Prkca* showed high expression levels in both human and mouse astrocytes (Figure 11D).

In microglia, the common neuropathic/inflammatory pain gene *GRM5/Grm5* was highly expressed in mice but slightly in humans (Figure 12A and B). The neuropathic pain-related genes *P2RX4/P2rx4* (purinergic receptor P2X 4, which functions as a ligand-gated ion channel with high calcium permeability), *HMGB1/Hmgb1*, *MAPK14/Mapk14* (mitogen-activated protein kinase 14), *PIK3CB/Pik3cb* (phosphatidylinositol-4,5-bisphosphate 3-kinase catalytic subunit beta, associated with immune responses after injury or infection), and *CRK/Crk* (CRK proto-oncogene, which encodes a member of an adapter protein family that binds to several tyrosine-phosphorylated proteins) were highly expressed in humans, but slightly in mice (Figure 12A). As in astrocytes, the inflammatory pain-related gene *PRKCA/Prkca* was highly expressed in both human and mouse microglia (Figure 12B).

In oligodendrocytes, the common neuropathic/inflammatory pain-related genes *GRM5/Grm5* were highly expressed in mice but slightly in humans (Figure 12C and D). *GRAP2/Grap2* (GRB2 related adaptor protein 2, involved in leukocyte-specific protein-tyrosine kinase signaling) was highly expressed in human Cluster 5 but slightly in mice (Figure 11C and D). *AIMP2/Aimp2* (aminoacyl-tRNA synthetase complex interacting multifunctional protein 2, part of the aminoacyl-tRNA synthetase complex) was highly expressed in human Cluster 4 but slightly in mouse (Figure 11C and D). As in microglia and astrocytes, the inflammatory pain-related genes *PRKCA/Prkca* were highly expressed in both human and mouse oligodendrocytes (Figure 11D).

We also examined the transcriptional profiles of the risk genes of other spinal cord diseases in human and mouse glial cells, including one type of motor neuron disease (ALS), two types of inflammatory diseases (multiple sclerosis and herpes zoster), and three types of developmental abnormalities (syringomyelia, diastematomyelia, and hereditary ataxias) (Supplementary Figures 4-7).

### Functional assignment for glial clusters

A Gene Ontology (GO) term analysis was performed to explore the functional properties of the enriched genes in glial cells and their subtypes; its results showed several common biological processes between humans and mice. The enriched genes in human and mouse glial cells were mainly associated with nervous system development, including neuronal migration, neuron projection development, axon guidance, axonogenesis, axon regeneration, dendrite morphogenesis, myelination; synaptic transmission, including chemical synaptic transmission; neurotransmitter reuptake, synapse assembly (Supplementary Figure 8). Detailed information on the GO term analysis for glial cell types and subtypes is shown in Supplementary Figures 9-12.

## Discussion

Despite the emerging evidence from animal models of the existence of distinct transcriptional profiles in glial cells [3, 10, 12, 13], little information is known about the heterogeneity within glial cells from human tissues. To the best of our knowledge, this is the first study to elucidate the cellular and molecular complexity of glial cells, including SGCs in the DRG and astrocytes, microglia, and oligodendrocytes in the spinal cord using 10× Genomics snRNA-seq. Furthermore, we compared human transcriptomic data with those from mice to explore conservation and divergence across species. Additionally, we examined the expression profiles of DRG and spinal cord disease-related genes in human and mouse glial cells.

The biological function of SGCs has been less studied than that of their CNS counterparts, astrocytes. SGCs envelop neurons in the PNS, mainly including the sensory, parasympathetic, and sympathetic ganglia [19]. SGCs activity is reportedly altered after neuronal injury in peripheral sensory ganglia, such as the DRG and trigeminal ganglia [20–23]. Despite the clear evidence that SGCs play an important role in the PNS, limited information is available regarding their cellular and molecular heterogeneity. van Weperen et al. performed scRNA-seq of mouse stellate ganglia and identified six distinct subtypes based on transcriptomic profiles [3]. However, our findings showed that SGCs cannot be completely distinguished into different subtypes either in human and mouse DRG, indicating low cellular heterogeneity within DRG SGCs.

Microglia are critically involved in many physiological and pathological conditions, such as sensation, pain, and neurodegeneration [1, 6, 24]. Previous studies have identified several distinct transcriptomic subtypes in brain microglia at single-cell resolution [9, 13, 14]. Using scRNA-seq, Geirsdottir et al. characterized the conservation and divergence of brain microglia across species. Interestingly, human brain microglia show significant heterogeneity compared with microglia in other mammals [9]. Similarly, we found that spinal microglia are also highly heterogeneous between humans and mice, as indicated by the slight overlap of transcriptomic profiles in the Uniform Manifold Approximation and Projection (UMAP). Likewise, SGCs, astrocytes, and oligodendrocytes exhibit substantial heterogeneity between humans and mice. In addition, the proportion of glial cell subtypes was different between humans and mice. Microglia represent 17.6% of all spinal cells in humans and only 1.2% in mice [17], whereas oligodendrocytes represent 52.4% of all spinal cells in humans and only 15.4% in mice [17]. These findings suggest that unlike peripheral DRG neurons [25], which have great similarity between humans and mice, glial cells show high divergence through evolution.

Rodent animal models are critical for understanding the molecular mechanisms underlying many physiological and pathological conditions. However, successful translation from animal models of neurodegenerative and chronic pain to human clinical trials is far from adequate [26, 27]. A previous study compared the gene expression of neurodegenerative disease susceptibility genes in human microglia with those of other mammalian species [9]. They observed significant expression changes in susceptibility genes for Alzheimer’s and Parkinson’s disease in humans compared with in rodents. In the present study, to better understand the conservation and divergence of microglial genes associated with common DRG and spinal cord diseases, we compared the transcriptional data of human microglia with those of mouse models in two common types of chronic pain, namely, neuropathic and inflammatory pain. As a result, significant heterogeneity in the expression levels of several genes was identified between humans and mice. Previous study showed that *Cxcr4* were widely expressed in rodents DRG SGCs [28] and CXCR4 signaling was associated with nerve injury-induced neuropathic pain [29]. In our study, *Cxcr4* was also highly expressed in mouse SGCs, however, it was almost absent in humans. In addition to *Cxcr4*, many disease-related genes were found highly expressed in mouse glial cells but almost absent in humans, such as *Calca*, *Gja1*, and *Kcnj10*. Moreover, several classical genes associated with nociceptive signaling, such as *Scn9a*, *Scn10a*, *Maf*, and *Kcna2*, were also highly expressed in mouse DRG SGCs, but not present in humans. These heterogeneous gene expression patterns between humans and mice, both in classical and disease marker genes, may partially explain the failure in the transition from animal models to clinical trials. Therefore, this work provides an important background to investigate the expression profiles of molecular targets of interest in glial cells for DRG and spinal cord disease.

This study has several limitations. First, the functional heterogeneity of distinct glial subtypes was not explored. Second, due to the small sample size, we did not examine the existence of sex-related heterogeneity of glial cells. Finally, the results of our comparison between humans and mice may be affected by the quality control in previous studies [16, 17].

In summary, our study provides a map of the complex cellular composition and transcriptomic atlas of glial cells in the human DRG and spinal cord, and provides an essential resource of conserved and divergent expression profiles in glial cells between humans and mice.

## Materials and Methods

### Human samples

This study has been approved by the Ethical Committee of the Affiliated Hospital of Zunyi Medical University (Approval No. KLL-2020-273, May 19, 2021). We have registered this study in the Chinese Clinical Trial Registry on June 20, 2021 (ChiCTR2100047511). Written informed consent was signed before patient enrollment. The study protocol was in consistent with the ethical and legal guidelines. The lumbar L3-5 DRGs and lumbar enlargements of spinal cord were acutely isolated from adult brain-dead human donors (two males, 38 and 46 years old; one female, 35 years old) withing 90 min of cross-clamp. These donors had not been diagnosed with acute/chronic low back or lower limb pain, two were dead from cerebral hernia and one was dead from intraventricular hemorrhage. All procedures were performed by the same surgeons from department of orthopedic surgery. Samples were immediately cleaned and frozen in liquid nitrogen for further use.

### Isolation of nuclei

The nucleus was isolated using a Nucleus Isolation Kit (catalogue no. 52009- 10, SHBIO, China) according to the manufacturer’s protocols. Briefly, frozen human samples were thawed on ice, minced, and homogenized in cold 1% BSA in lysis buffer. The lysates were filtered through a 40-μm cell strainer, followed by a centrifugation (500 × g, 5 min) at 4°C. Then, the pellets were resuspended in the lysis buffer after removing the supernatant. PBS were added and centrifuged (3000 × g, 20 min) at 4°C. The pellets were then filtered through a 40-μm cell strainer, centrifuged (500 g, 5 min) at 4°C, and resuspended twice in the nuclease-free BSA. Finally, nuclei were stained by trypan blue and counted using a dual-fluorescence cell counter.

### cDNA synthesis

The nuclei suspension was loaded onto a Chromium single cell controller (10x Genomics) to produce single-nucleus gel beads in the emulsion (GEM) using single cell 3’ Library and Gel Bead Kit V3.1 (10x Genomics, 1000075) and Chromium Single Cell B Chip Kit (10x Genomics, 1000074) according to the manufacturer’s instructions. After the captured nucleus was lysed, the released mRNA was barcoded using reverse transcription in individual GEM. cDNA was generated by reverse transcription using a S1000TM Touch Thermal Cycler (Bio-Rad, 53°C for 45 min, 85°C for 5 min, and 4°C until further use). The cDNA was then amplified and their quality was determined using an Agilent 4200 (CapitalBio Technology, Beijing).

### 10x Genomics library preparation and sequencing

The 10x snRNA-seq library was established using Single Cell 3’ Library and Gel Bead Kit V3.1 according to the manufacturer’s instructions. The library was sequenced using a Novaseq 6000 sequencing platform (Illumina) with a depth of at least 30,000 reads per nucleus with a paired-end 150 bp (PE150) reading strategy (CapitalBio Technology, Beijing).

### Single-nucleus transcriptomic data analysis Data pre-processing

We first processed 10X genomics raw data by the Cell Ranger Single-Cell Software Suite (release 5.0.1), including using cellranger mkfastq to demultiplexes raw base call files into FASTQ files and then using cellranger count to preform alignment, filtering, barcode counting, and UMI counting. The reads were aligned to the hg19 reference genome using a pre-built annotation package download from the 10X Genomics website.

### Quality control

We performed quality control to remove low-quality cells, empty droplets or cell doublets. Low-quality cells or empty droplets often contain very few genes or exhibit extensive mitochondrial contamination, whereas cell doublets may exhibit an aberrantly high gene count. Moreover, we can detect contamination with low complexity cells like red blood cells that show a less complex RNA species. Specifically, cells were filtered out if the gene number was less than 200 or more than 8,000, UMI counts was less than 500, cell complexity was less than 0.8, or if the mitochondrial gene ratio was more than 25%.

### Normalization and integration

Seurat package was used to normalize and scale the single-nucleus gene expression data. Data was first normalized by ‘‘Normalize Data’’ function with setting normalization method as ‘‘Log Normalize.’’ In detail, the expression of gene A in cell B was determined by the UMI count of gene A divided by the total number of UMI of the cell B, followed by multiplying 10,000 for the normalization and the log-transformed counts were then computed with base as 2. Top 2,000 highly variable genes (HVGs) were detected by “Find Variable Features” function with setting selection method as “vst”. We then removed the uninteresting sources of variation by regressing out cell-cell variation within gene expression driven by batch, the number of detected UMI, mitochondrial gene expression, and ribosomal gene expression, which was implemented by ‘‘Scale Data’’ function. Finally, the corrected expression matrix was used as an input for further analysis.

### Dimension reduction, cell clustering and annotation

“Run PCA’’ function in the Seurat package was used to perform the principal component analysis (PCA) on the single-cell expression matrix with genes restricted to HVGs. To integrate cells into a shared space from different batches for unsupervised clustering, harmony algorithm was used to integrate two batches, which was implemented by “Run Harmony” function. ‘‘Find Clusters’’ function in the Seurat package was then used to conduct the cell clustering analysis through embedding cells into a graph structure in harmony space. The clustering results were visualized using UMAP. Multiple cell type-specific/enriched marker genes that have been previously described in the literature were used to determine cell-type identity. These include *FABP7* and *APOE* for SGCs; *ATP1A2*, *AQP4*, *GJA1*, and *SLC1A2* for Astrocytes; *PTPRC*, *CTSS*, and *ITGAM* for Microglia; *MBP*, *MOBP*, *MOG*, and *PLP1* for oligodendrocytes.

### Differential expression analysis

Differential gene expression analysis was performed using the ‘Find Markers’ function, which performs differential expression based on the non-parameteric Wilcox rank sum test for two annotated cell groups. The marker genes were identified using the ‘Find All Markers’ function in Seurat with settings on genes with at least 0.25 increasing logFC upregulation, comparing to the remaining cell clusters.

### Enrichment analysis

GO enrichment was conducted using R package with default settings. GO terms with an adjusted p-value of less than 0.05 that calculated by the hypergeometric test followed by the Benjamini-Hochberg method were defined as significantly enriched terms. The top 10 enriched terms were visualized.

### Risk genes of diseases

The risk genes of spinal cord diseases were identified using DisGeNET (https://www.disgenet.org), which contains one of the largest publicly available collections of genes associated with human diseases. The current version of DisGeNET (v7.0) contains 1,134,942 gene-disease associations (GDAs), between 21,671 genes and 30,170 diseases, disorders, traits, and clinical or abnormal human phenotypes. From the summary of GDAs, the top 50 risk genes ordered by the number of PMIDs were selected.

### Comparison to RNA-seq data sets from mouse

For analysis of the mouse, a random subset of data from sn-RNA sequencing of DRGs and spinal cord from wild type mice was extracted from data deposited by Renthal et al. [16] and Sathyamurthy et al. [17], respectively.

## Acknowledgments

Thanks to Hongjun Chen and Jin Li from Affiliated Hospital of Zunyi Medical University for providing assistance in sample collection. Thanks to Yang Yang from Basebio for providing data analysis assistance. This work was supported by the National Key Research and Development Program of China (Project No. 2020YFC2008400 and 2020YFC2008402) (To Cheng Zhou); Grant No. 81974164 (To Cheng Zhou) and Grant No. 82001183 (To Mengchan Ou) from the National Natural Science Foundation of China (Beijing, China); and Grant No. 2021M692276 (To Donghang Zhang) from China Postdoctoral Science Foundation.

## Author Contributions

C.Z. and D.Z. conceived the study. C.Z., D.Z., T.Z., and J.L. designed the study. D.Z., Y.W., Y.Y., M.O., Y.C., and J.S. performed the experiments and analyzed the data. C.Z., J. Liu, T.Z., and D.Z. wrote the manuscript.

## Data and code availability

The accession number for the raw sequencing data and processed data reported in this paper is GEO: GSE 189501.

## Lead contact

Further information and requests for resources and reagents should be directed to and will be fulfilled by the lead contact, Cheng Zhou.

## Supplementary figures and legends

**Supplementary Figure 1.**
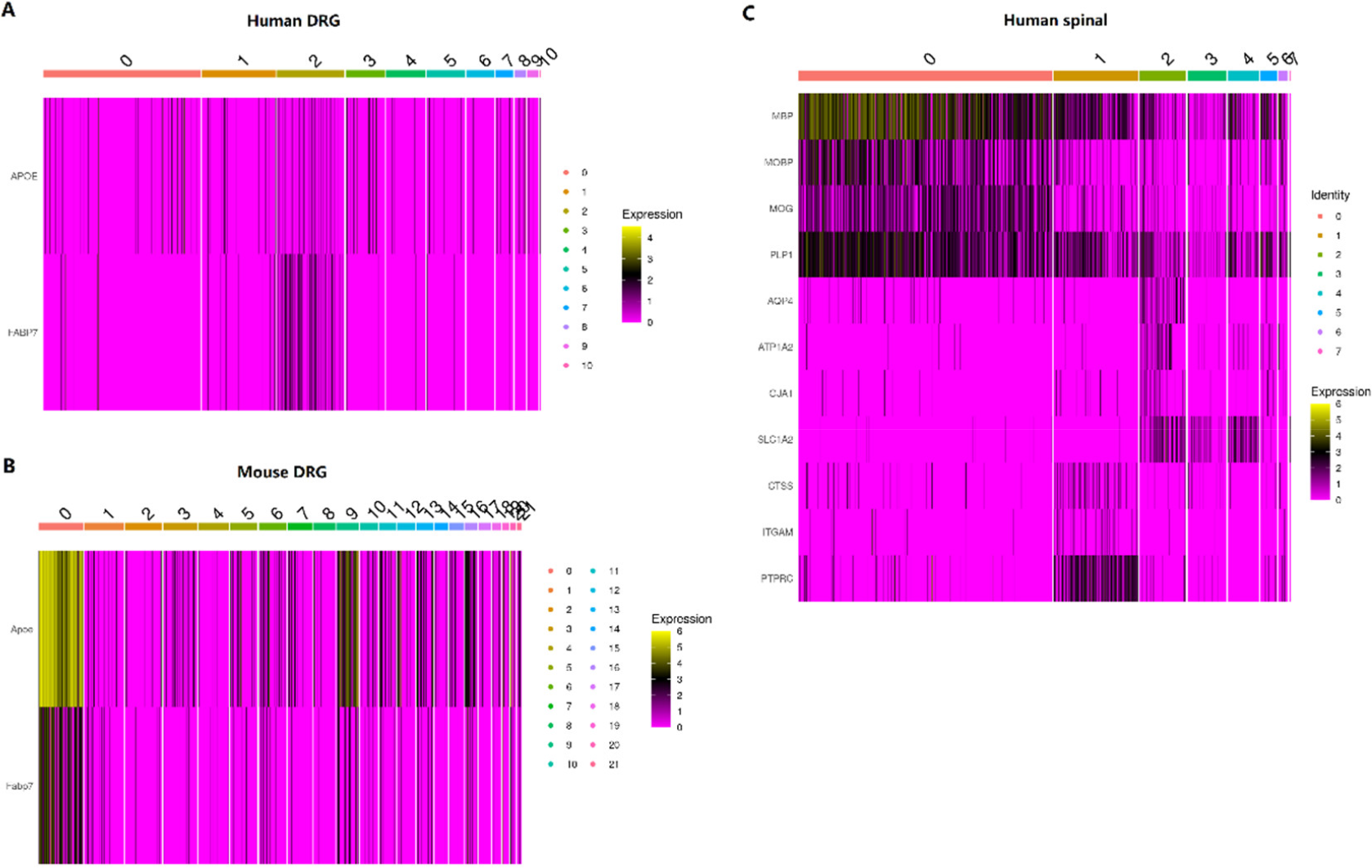
(A-B) Heatmap showing the distribution of expression levels of marker genes of human (A) and mouse (B) DRG SGCs across all seven cell types. (C) Heatmap showing the distribution of expression levels of marker genes of human spinal astrocytes, microglia, and oligodendrocytes across all cell types.

**Supplementary Figure 2.**
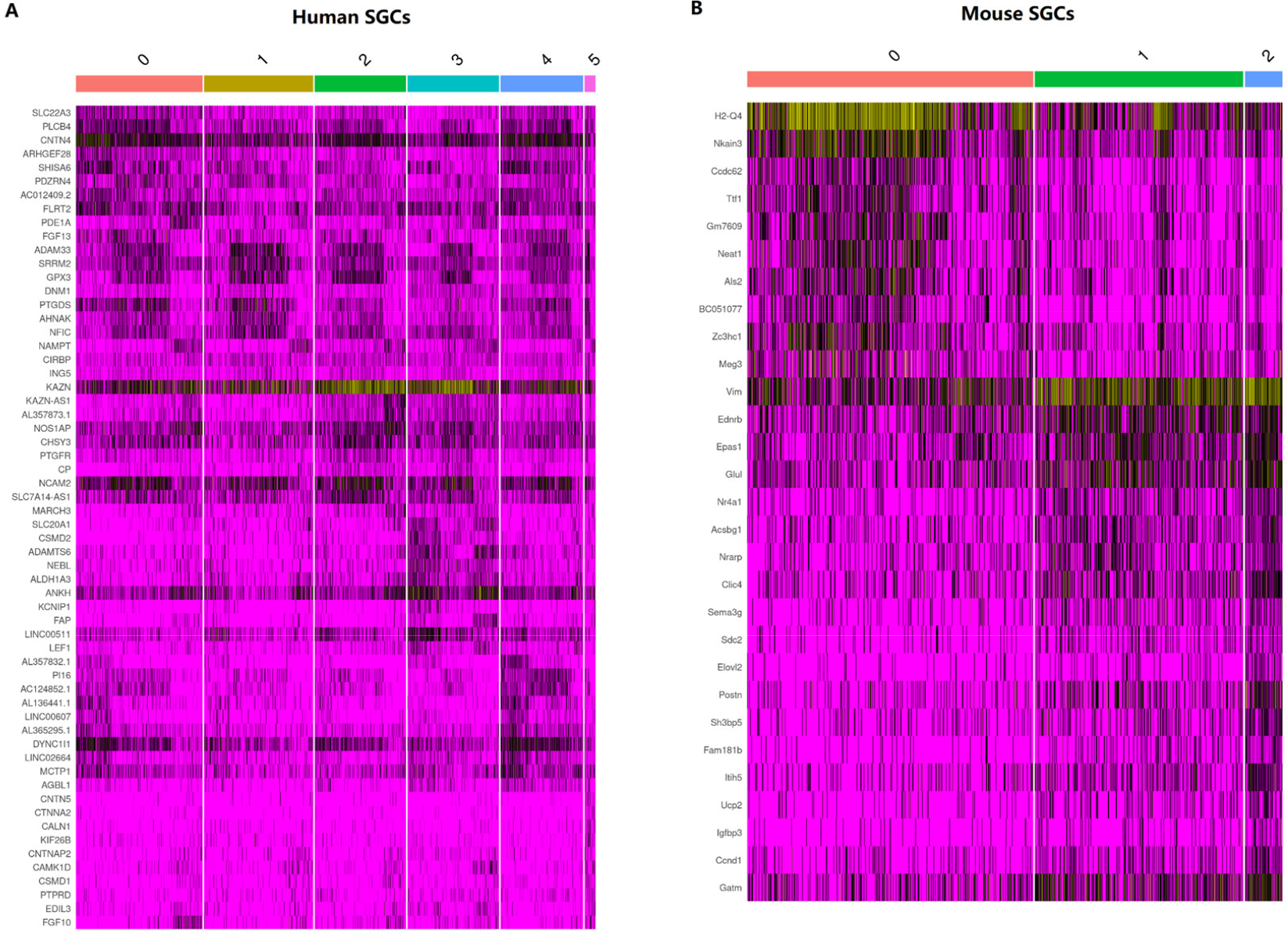
(A-B) Heatmap showing the expression of the top ten most differentially expressed genes across all the subclusters of DRG SGCs in human (A) and mouse (B). SGCs, satellite glial cells.

**Supplementary Figure 3.**
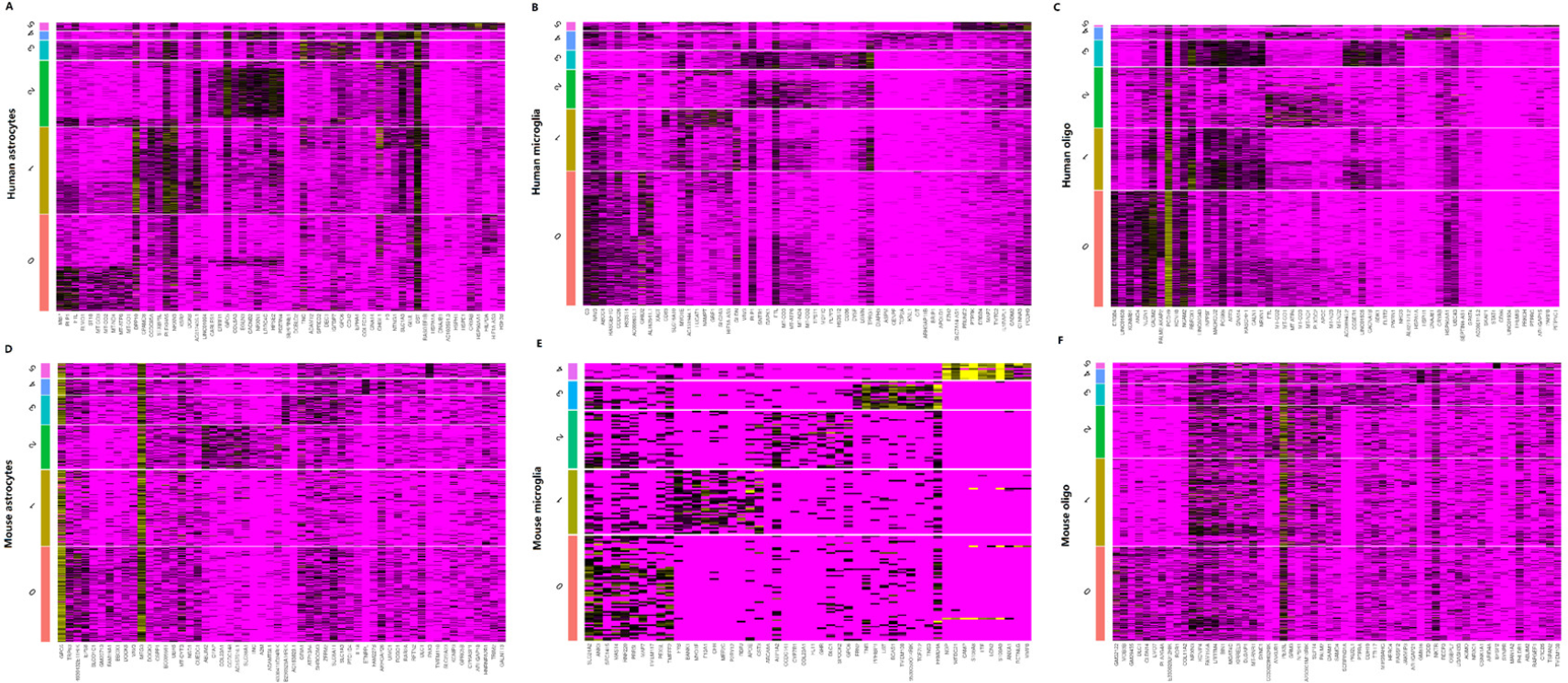
(A-C) Heatmap showing the expression of the top ten most differentially expressed genes across all the subclusters of spinal astrocytes (A), microglia (B), and oligodendrocytes (C) in humans. (D-F) Heatmap showing the expression of the top ten most differentially expressed genes across all the subclusters of spinal astrocytes (D), microglia (E), and oligodendrocytes (F) in mice. Oligo, oligodendrocytes.

**Supplementary Figure 4.**
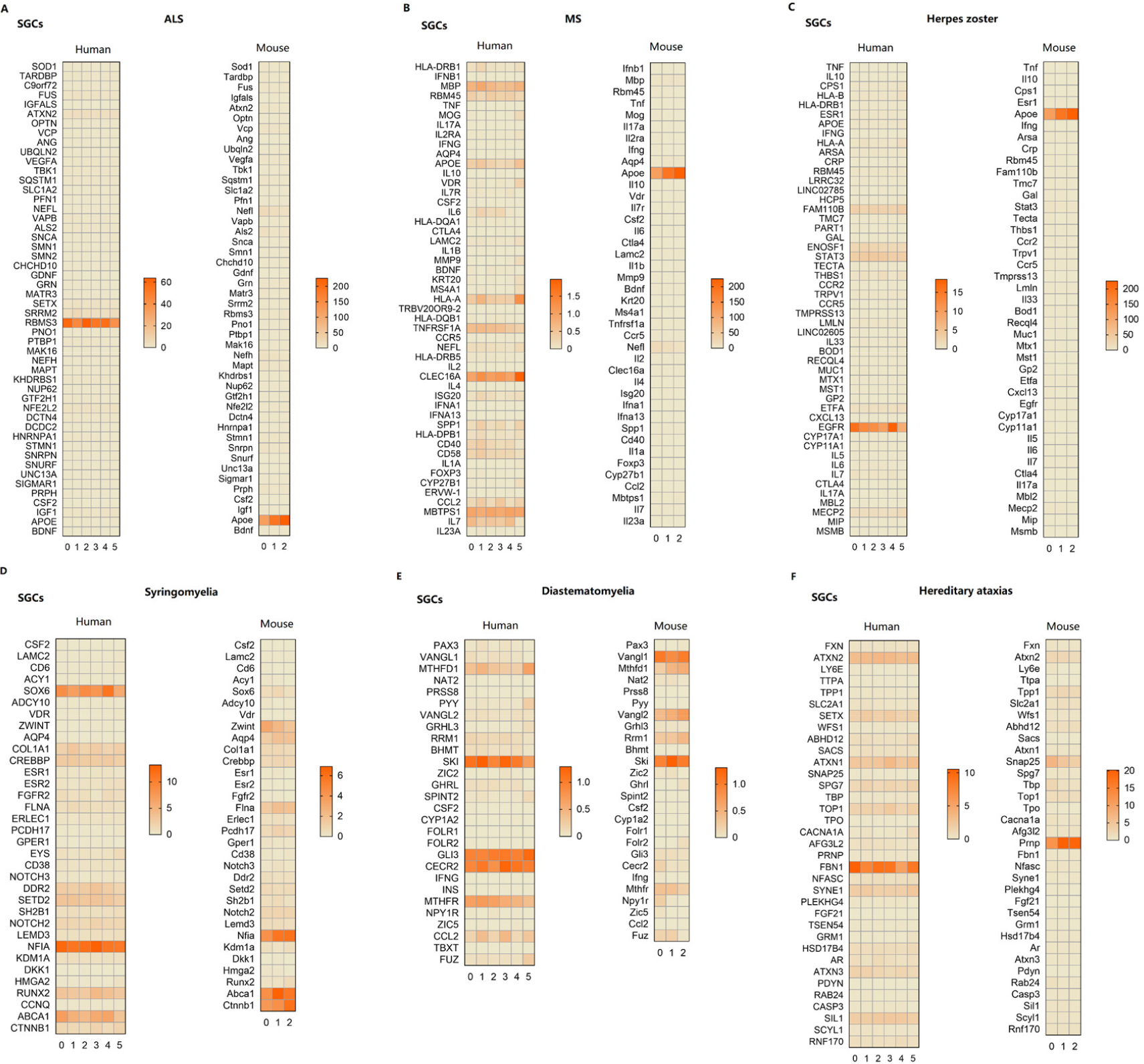
(A-G) Expression profiles of risk genes for ALS (A), MS (B), Herpes zoster (C), Syringomyelia (D), Diastematomyelia (E), and Hereditary ataxias (F) in DRG SGCs of humans and mice. SGCs, satellite glial cells; ALS, Amyotrophic lateral sclerosis; MS, Multiple sclerosis.

**Supplementary Figure 5.**
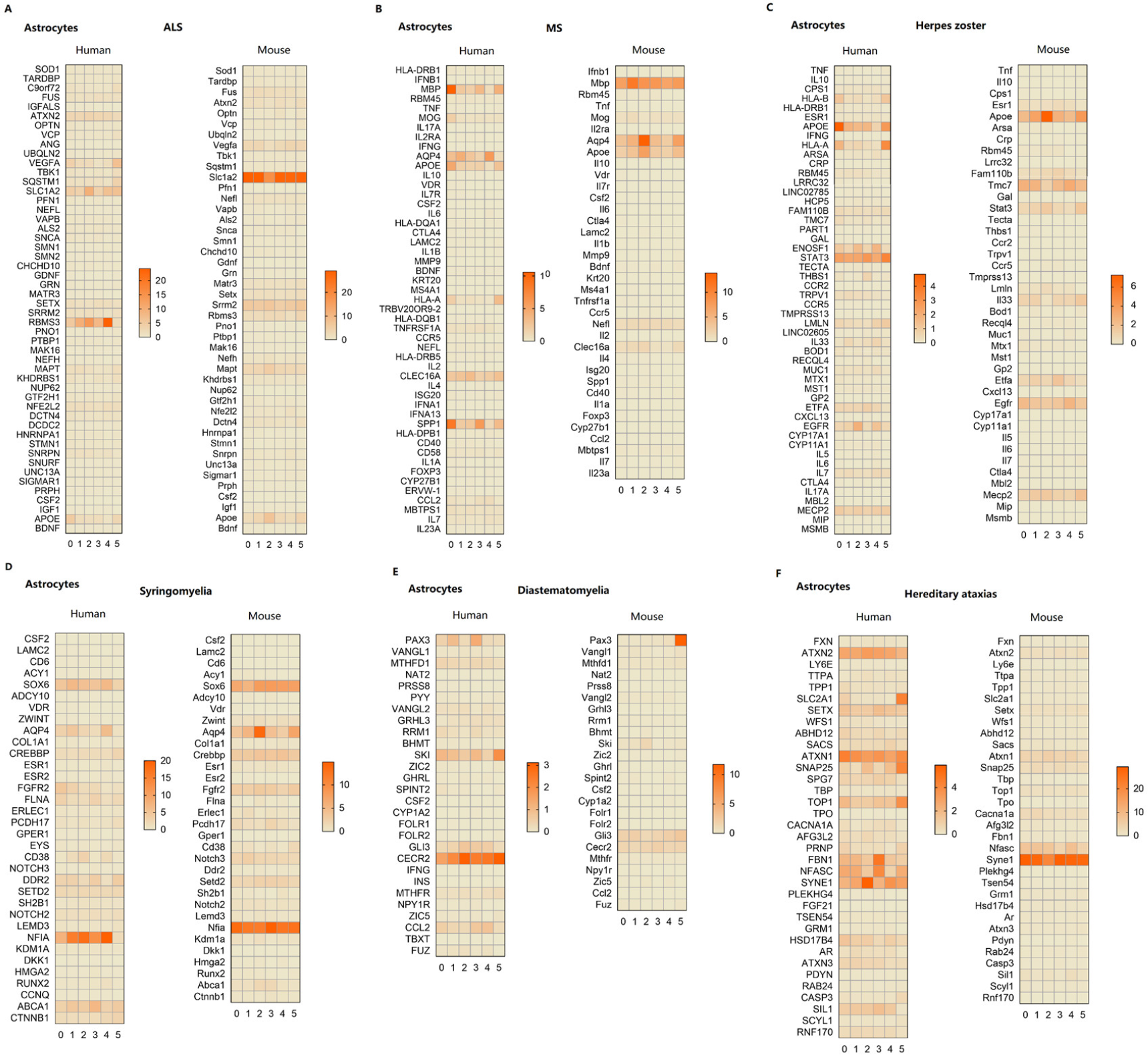
(A-G) Expression profiles of risk genes for ALS (A), MS (B), Herpes zoster (C), Syringomyelia (D), Diastematomyelia (E), and Hereditary ataxias (F) in astrocytes of humans and mice. ALS, Amyotrophic lateral sclerosis; MS, Multiple sclerosis.

**Supplementary Figure 6.**
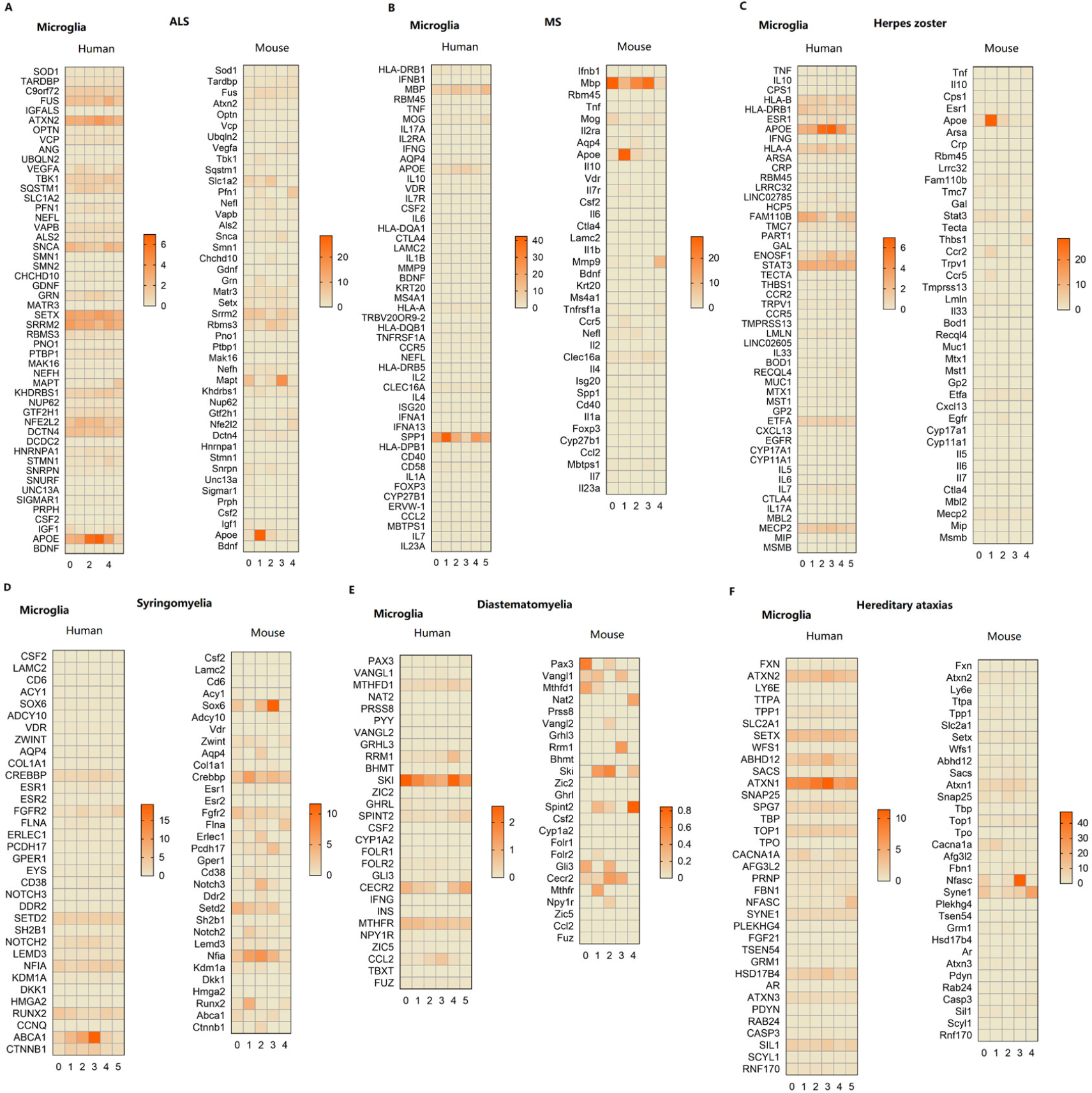
(A-G) Expression profiles of risk genes for ALS (A), MS (B), Herpes zoster (C), Syringomyelia (D), Diastematomyelia (E), and Hereditary ataxias (F) in microglia of humans and mice. ALS, Amyotrophic lateral sclerosis; MS, Multiple sclerosis.

**Supplementary Figure 7.**
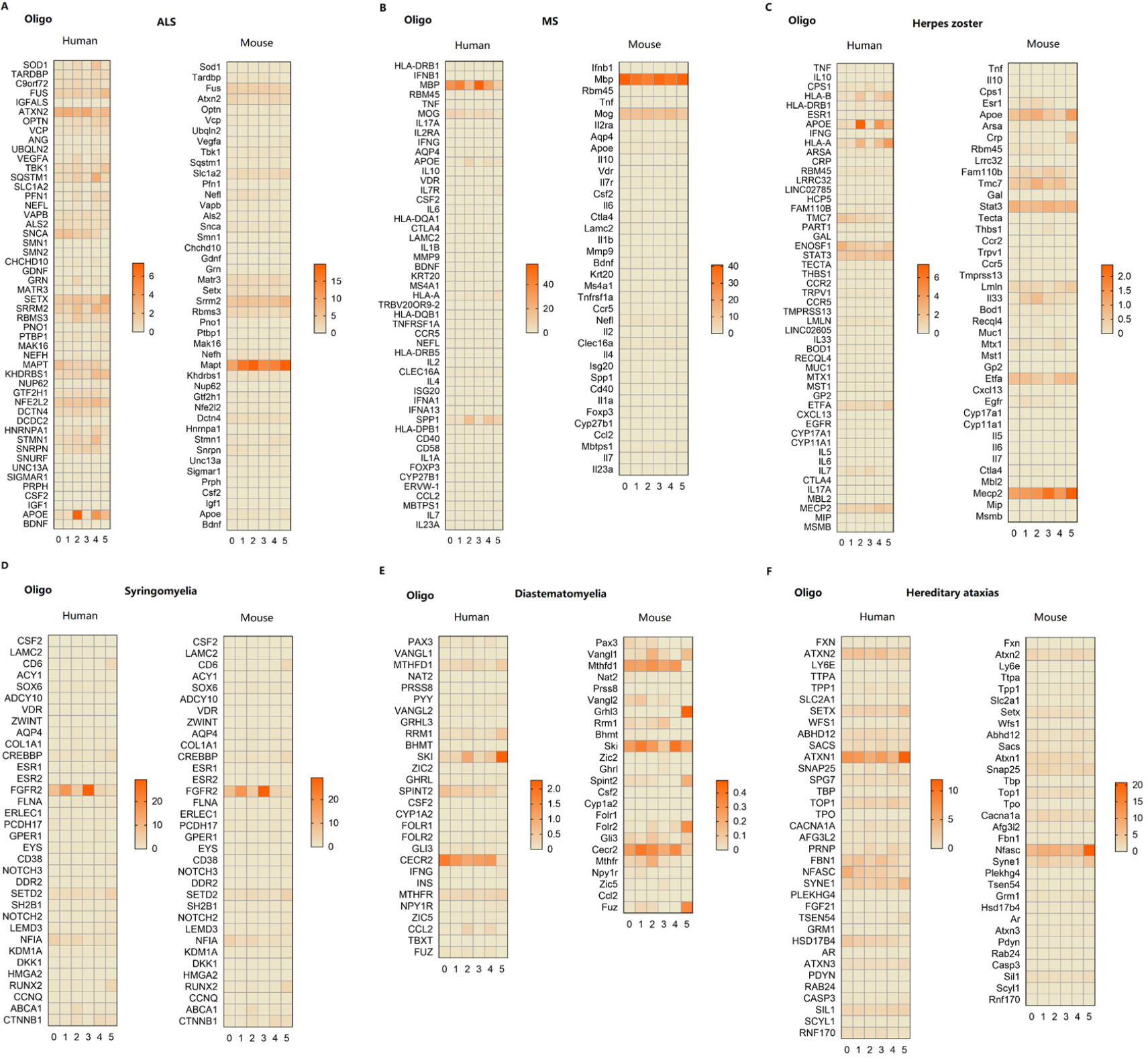
(A-G) Expression profiles of risk genes for ALS (A), MS (B), Herpes zoster (C), Syringomyelia (D), Diastematomyelia (E), Hereditary ataxias (F) in oligodendrocytes of humans and mice. oligo, oligodendrocytes; ALS, Amyotrophic lateral sclerosis; MS, Multiple sclerosis.

**Supplementary Figure 8.**
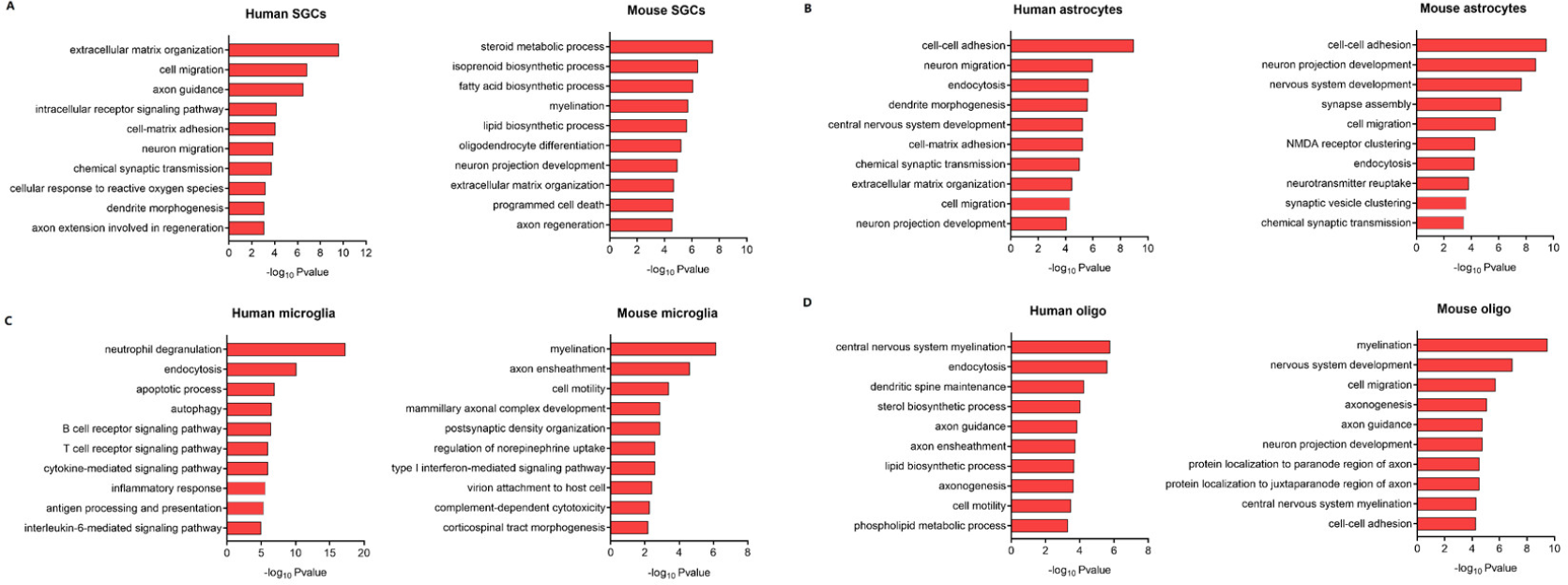
(A-D) Summarized GO terms for the enriched genes in SGCs (A), astrocytes (B), microglia (C), and oligodendrocytes (D) of humans and mice. SGCs, satellite glial cells; oligo, oligodendrocytes.

**Supplementary Figure 9.**
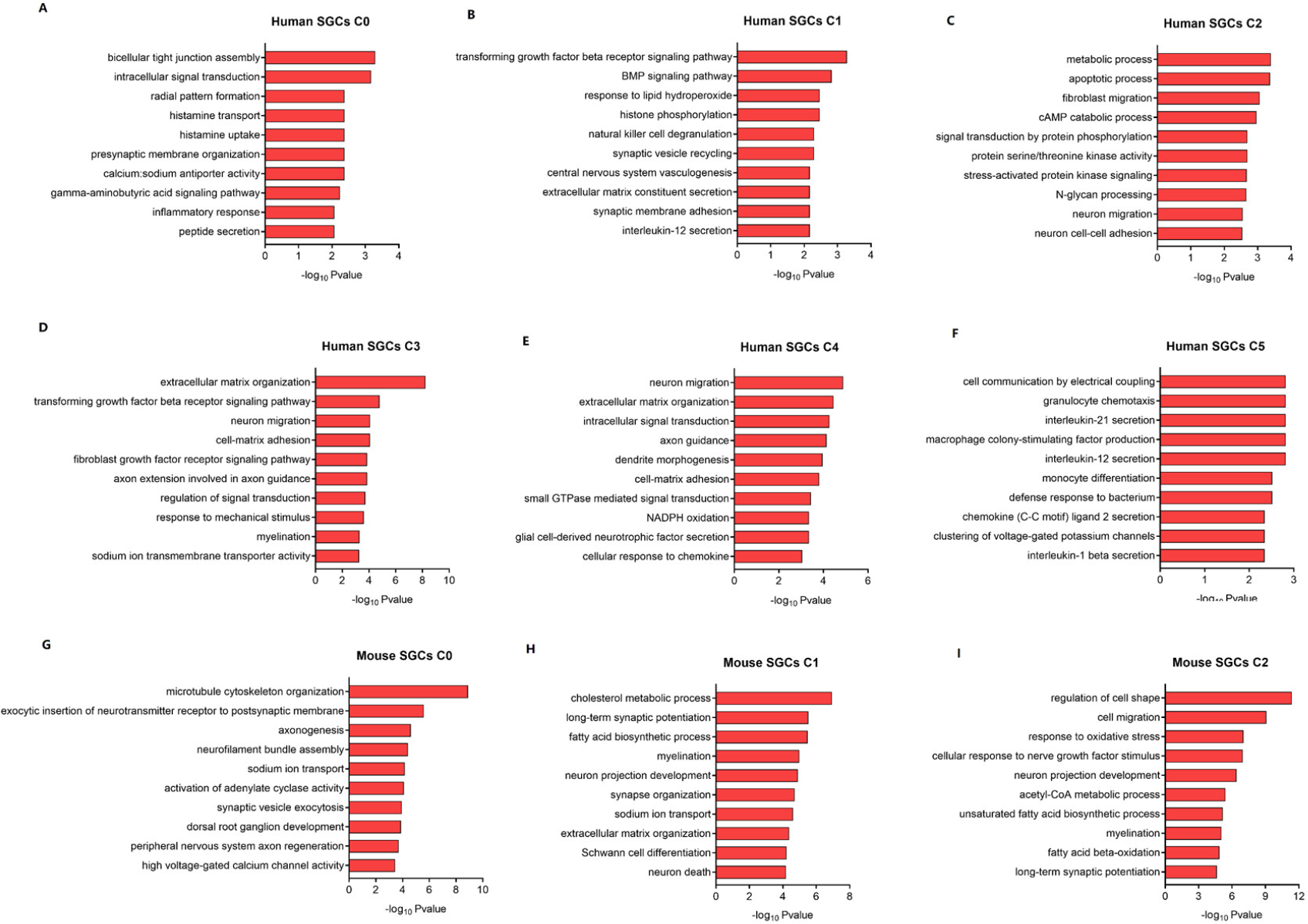
(A-F) Summarized GO terms for the enriched genes in each subtype of human SGCs. (G-I) Summarized GO terms for the enriched genes in each subtype of mouse SGCs. SGCs, satellite glial cells.

**Supplementary Figure 10.**
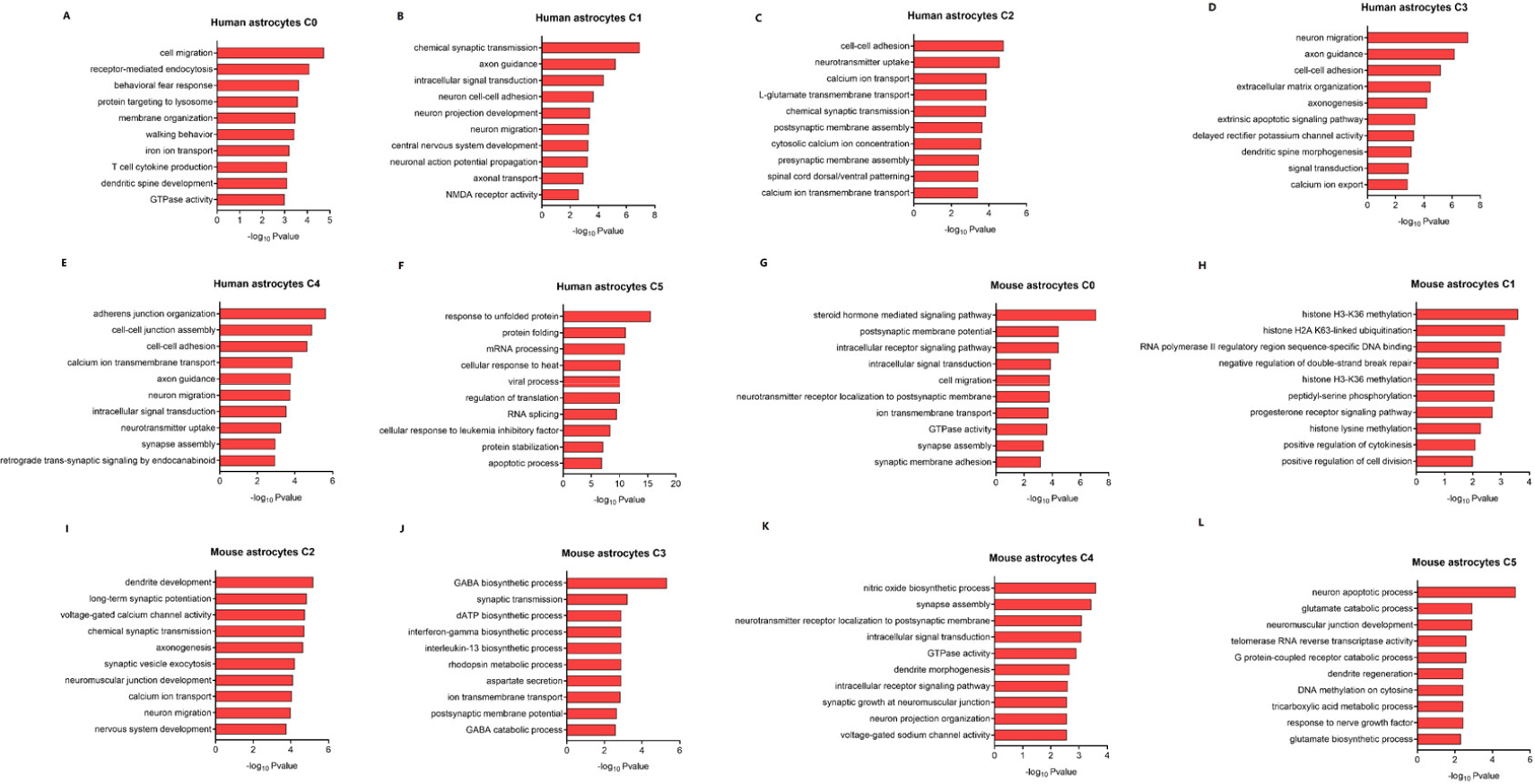
(A-F) Summarized GO terms for the enriched genes in each subtype of human astrocytes. (G-L) Summarized GO terms for the enriched genes in each subtype of mouse astrocytes.

**Supplementary Figure 11.**
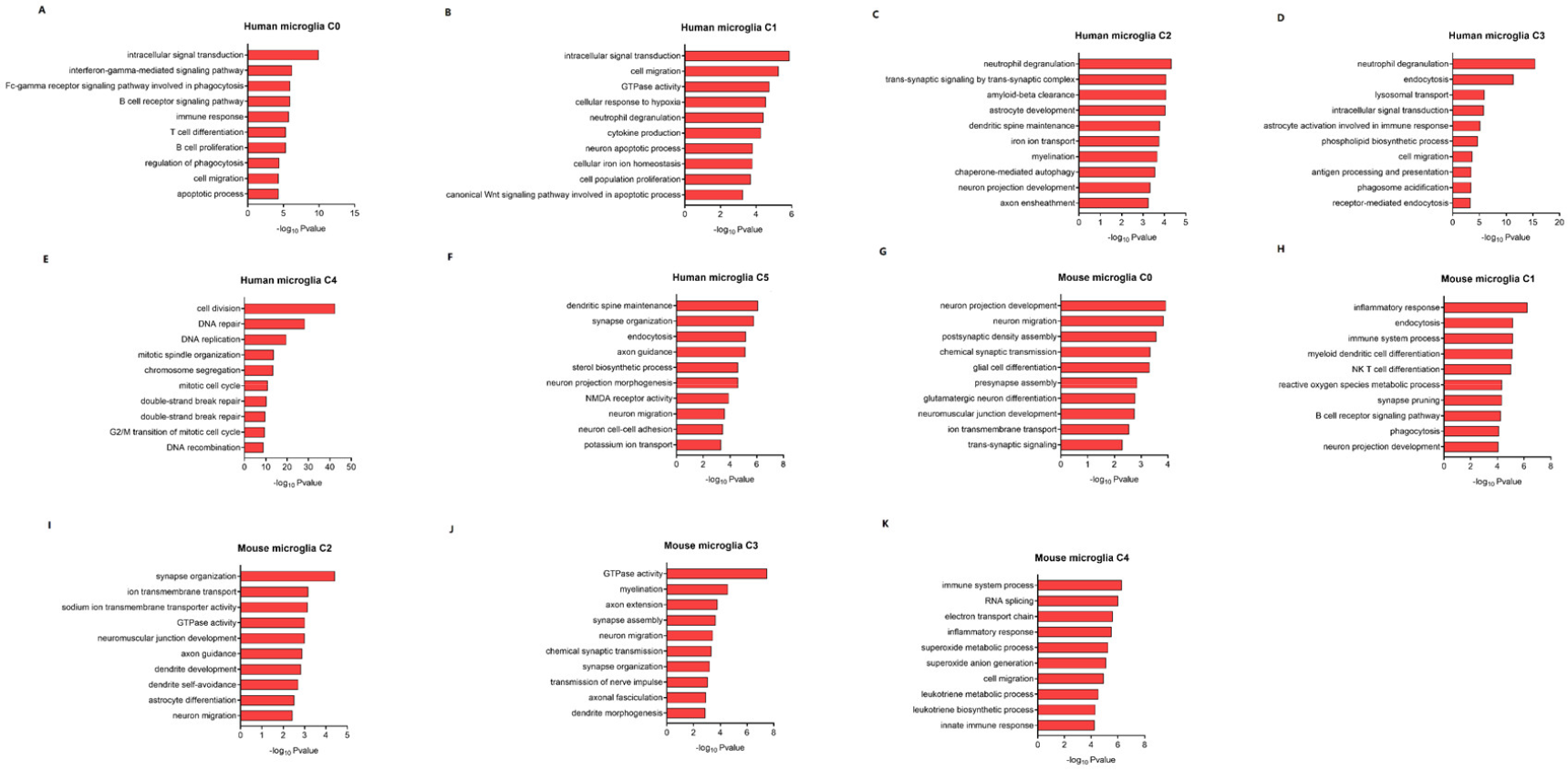
(A-F) Summarized GO terms for the enriched genes in each subtype of human microglia. (G-K) Summarized GO terms for the enriched genes in each subtype of mouse microglia.

**Supplementary Figure 12.**
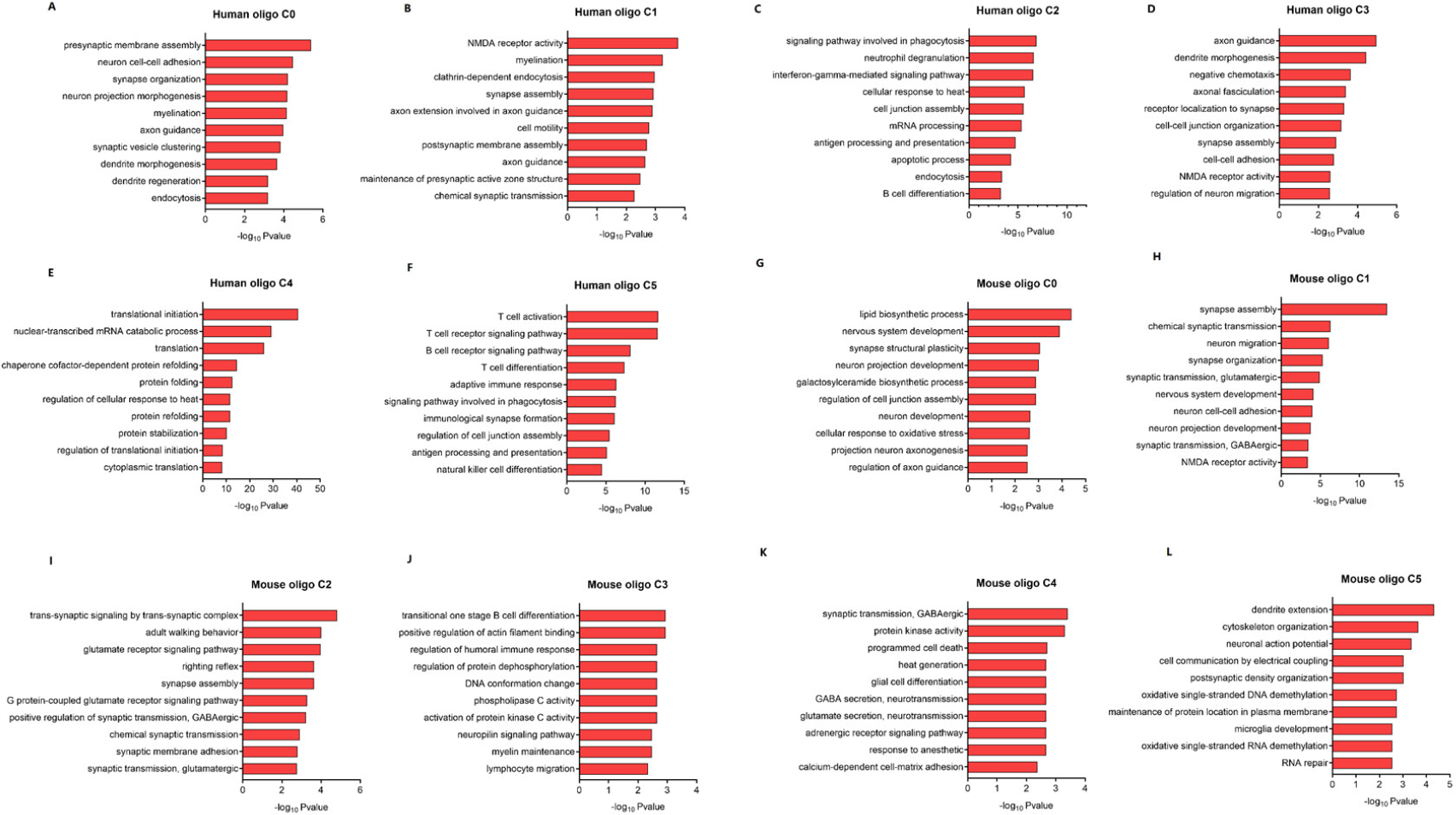
(A-F) Summarized GO terms for the enriched genes in each subtype of human oligodendrocytes. (G-K) Summarized GO terms for the enriched genes in each subtype of mouse oligodendrocytes. SGCs, satellite glial cells; oligo, oligodendrocytes.

